# Hypoxia-inducible lipid droplet-associated interacts with DGAT1 and promotes lipid storage in hepatocytes

**DOI:** 10.1101/2020.02.26.966374

**Authors:** Montserrat A. de la Rosa Rodriguez, Anne Gemmink, Michel van Weeghel, Marie Louise Aoun, Christina Warnecke, Rajat Singh, Jan Willem Borst, Sander Kersten

## Abstract

Lipid droplets (LD) are dynamic organelles that can expand and shrink, driven by fluctuations in the rate of triglyceride synthesis and degradation. Triglyceride synthesis, storage in LD, and degradation are governed by a complex set of LD-associated proteins. One of these LD-associated proteins, hypoxia-inducible lipid droplet-associated (HILPDA), was found to impair LD breakdown by inhibiting adipose triglyceride lipase. Here we characterized the physiological role and mechanism of action of HILPDA in hepatocytes. Expression of HILPDA was induced by fatty acids in several hepatoma cell lines. Hepatocyte-specific deficiency of HILPDA in mice modestly but significantly reduced hepatic triglycerides in mice with non-alcoholic fatty liver disease. Similarly, deficiency of HILPDA in mouse precision-cut liver slices and primary hepatocytes reduced lipid storage and accumulation of fluorescently-labelled fatty acids in LD, respectively, which was independent of adipose triglyceride lipase. Fluorescence microscopy showed that HILPDA partly colocalizes with LD and with the endoplasmic reticulum, is especially abundant in perinuclear areas, and mainly associates with newly added fatty acids. Real-time fluorescence live-cell imaging further revealed that HILPDA preferentially localizes to LD that are being remodelled. Mechanistically, HILPDA overexpression increased lipid storage in human hepatoma cells concomitant with an increase in DGAT activity and DGAT1 protein levels. Finally, confocal microscopy and Förster resonance energy transfer-fluorescence lifetime imaging microscopy analysis indicated that HILPDA colocalizes and physically interacts with DGAT1. Overall, our data indicate that HILPDA physically interacts with DGAT1 and increases DGAT activity. These findings suggest a novel mechanism in hepatocytes that links elevated fatty acid levels to stimulation of triglyceride synthesis and storage.

## INTRODUCTION

Fatty acids are an important fuel for many cell types. When the supply of fatty acids exceeds the demand for oxidation, excess fatty acids can be stockpiled by converting them to triglycerides. Triglycerides are synthesized in the endoplasmic reticulum and are stored in specialized organelles called lipid droplets (LD) (18). With the exception of adipocytes, most cell types have tiny LD that collectively only take up a very small portion of the total cell volume. However, in certain pathological conditions, LD may become enlarged and occupy considerable cell volume, potentially interfering with important cellular functions (17).

The liver plays a central role in the regulation of lipid metabolism. Under conditions of obesity and insulin resistance, storage of lipids in the liver is often elevated (40). A chronic increase in intra-hepatic fat is referred to as steatosis and is a key feature of non-alcoholic fatty liver disease (NALFD) (29). In many high-income countries, NAFLD has become the most common liver disorder and a growing clinical concern (29).

Fatty acids in hepatocytes can originate from several different sources: from triglycerides taken up as chylomicron-remnants, from endogenous synthesis (de novo lipogenesis), and from circulating non-esterified fatty acids released by adipose tissue (20). A major portion of the incoming fatty acids is oxidized to provide energy to hepatocytes. The remainder is esterified into triglycerides, part of which is incorporated and secreted in very low-density lipoprotein, and part of which is stored in LD in hepatocytes. Accordingly, excess storage of lipids in the liver can be due to changes in several metabolic pathways, including defective fatty acid oxidation, enhanced lipogenesis, impaired triglyceride secretion, and increased uptake of fatty acids from the circulation (21).

LD are dynamic organelles that can rapidly expand and shrink, driven by fluctuations in the rate of triglyceride synthesis and degradation (17). The synthesis of triglycerides, their storage in LD, and the subsequent breakdown of triglycerides into fatty acids are governed by a complex set of enzymes and LD-associated proteins. LD-associated proteins encompass a large group of proteins that physically and functionally interact with LD. According to proteomic profiling, the number of LD-associated proteins easily runs into hundreds (2, 6, 34, 39). This group includes lipid synthesis and degradation enzymes, proteins involved in membrane trafficking, lipid signaling proteins, and proteins involved in protein degradation (39). An important group of LD-associated proteins is the perilipin family, composed of PLIN1-PLIN5 (33). Other known LD-associated proteins include CIDEA, CIDEB, CIDEC, FITM1, FITM2, G0S2, and ABHD5 (7, 14, 43).

A relatively poorly characterized LD-associated protein is HILPDA. The first identification of HILPDA as LD-associated protein was in Hela cells, where its overexpression was found to increase intracellular lipid accumulation (15). HILPDA raised our attention when trying to identify novel target genes of the transcription factors PPARα and PPARγ in hepatocytes and adipocytes, respectively, and when screening for novel genes induced by fatty acids in peritoneal macrophages (11, 24, 35). In mouse liver, HILPDA overexpression via adeno-associated viral delivery raised intrahepatic triglyceride levels by approximately 4-fold, likely by suppressing very-low-density lipoprotein triglyceride secretion (24). Consistent with these data, deficiency of HILPDA in cultured hepatocytes lowered hepatic lipid accumulation, which was explained by a combination of decreased fatty acid uptake, increased fatty acid beta-oxidation, and increased triglyceride lipolysis (12). Somewhat surprisingly, hepatocyte-specific HILPDA deficiency did not influence liver triglyceride content in mice chronically fed a high fat diet (12).

Recently, we and others found that HILPDA is able to bind the intracellular triglyceride hydrolase ATGL and inhibit ATGL-mediated triglyceride hydrolysis (28, 35, 36, 44). Although the relatively low inhibitory capacity of HILPDA towards ATGL in cell-free systems raised questions about the physiological relevance of this interaction (28), studies in HILPDA-deficient macrophages and cancer cells firmly established the functional dependence between HILPDA and ATGL (35, 36, 44). Currently, very little is known about the molecular mechanism of action of HILPDA in hepatocytes. Accordingly, the present study was aimed at better characterizing the molecular role of HILPDA in hepatocytes.

## METHODS

### Mice experiments

*Hilpda*^flox/flox^ mice (Jackson Laboratories, Bar Harbor, ME; Hilpdatm1.1Nat, #017360) were acquired and crossed with C57Bl/6J mice for at least 5 generations. Thereafter, the *Hilpda*^flox/flox^ mice were crossed with Albumin-Cre transgenic mice (Jackson Laboratories, Bar Harbor, ME; B6.Cg-Speer6-ps1^Tg(Alb-cre)21Mgn^/J, #003574) to generate mice with hepatocyte-specific Cre-mediated deletion of *Hilpda* (*Hilpda*^Δhep^). Mice were group housed under normal light-dark cycles in temperature- and humidity-controlled specific pathogen-free conditions. Mice had ad libitum access to regular chow and water.

Male *Hilpda*^Δhep^ mice aged 4-5 months and their *Hilpda*^flox/flox^ littermates were fasted from 11:00h onwards and euthanized the next day between 11:00h and 12:00h (fasted group). Alternatively, mice were fasted from 11:00h onwards with reintroduction of chow the next morning at 07:30h, followed by euthanasia between 11:00h and 12:00h (refed group). The number of mice per group was 9-13.

Male *Hilpda*^Δhep^ mice aged 3-4 months and their *Hilpda*^flox/flox^ littermates were given a semi-purified low fat diet (10% kcal fat, A08051501) or high fat diet lacking choline and methionine (45% kcal fat, A06071309)(Research Diets, Inc. New Brunswick, NJ). During the dietary intervention, the mice were housed individually. After 11 weeks, mice were euthanized in the ad libitum fed state between 8:15h and 10.00h. The number of mice per group was 12.

Prior to euthanasia, mice were anaesthetised with isoflurane and blood was collected via orbital puncture in tubes containing EDTA (Sarstedt, Nümbrecht, Germany). Immediately thereafter, mice were euthanized by cervical dislocation, after which tissues were excised, weighed, and frozen in liquid nitrogen or prepared for histology. Frozen samples were stored at −80°C. Liver tissue was fixed in 4% formaldehyde solution in PBS. All animal experiments were approved by the local animal welfare committee of Wageningen University (AVD104002015236, 2016.W-0093.007 and 2016.W-0093.017). The experimenter was blinded to group assignments during all analyses.

### Plasma measurements

Blood collected in EDTA tubes (Sarstedt, Numbrecht, Germany) was spun down for 10 minutes at 2000 g at 4°C. Plasma was aliquoted and stored at −80°C until further measurements. The plasma concentration of various metabolites was determined used specialized kits: cholesterol (Liquicolor, Human GmbH, Wiesbaden, Germany), triglycerides (Liquicolor), glucose (Liquicolor), NEFAs (NEFA-HR set R1, R2 and standard, WAKO Diagnostics, Instruchemie, Delfzijl, The Netherlands), Alanine Transaminase Activity Assay Kit (Abcam ab105134), following the manufacturer’s instructions.

### Liver triglycerides

2% liver homogenates were prepared in a buffer (10 mM Tris, 2 mM EDTA and 0.25 M sucrose, pH 7.5) by homogenising in a Tissue Lyser II (Qiagen, Hilden, Germany). Liver triglyceride content was then quantified using Triglyceride liquicolor mono from HUMAN Diagnostics (Wiesbaden, Germany) according to the manufacturer’s instructions.

### Cell treatments and gene expression

Human HepG2, mouse Hepa 1-6 and rat Fao hepatoma cells at 75% confluency were incubated with a mixture of oleate and palmitate (ratio 2:1, total concentration 1.2 mM) coupled to FA-free Bovine Serum Albumin (BSA) (Roche Applied Sciences). All fatty acid stocks were initially reconstituted in absolute ethanol. Sub-stocks of fatty acids at 25 mM were prepared in filter-sterilised KOH at 70 mM. Fatty acids were diluted in DMEM containing 3% FA-free BSA to obtain the desired final concentrations. After treatment, cells were washed with ice-cold phosphate-buffered saline (PBS) (Lonza) and stored at −20°C for further analysis.

Total RNA was isolated using TRIzol® Reagent (Invitrogen, ThermoFisher Scientific). cDNA was synthesized from 500 ng RNA using the iScript cDNA kit (Bio-Rad Laboratories, Hercules, CA, USA) according to manufacturer’s instructions. Real time polymerase chain reaction (RT-PCR) was performed with the CFX96 or CFX384 Touch(tm) Real-Time detection system (Bio-Rad Laboratories), using a SensiMix(tm) (BioLine, London, UK) protocol for SYBR green reactions. Mouse 36b4 expression was used for normalization.

### Liver slices

Precision cut liver slices were prepared from *Hilpda*^Δhep^ and *Hilpda*^flox/flox^ mice as described previously (32). Briefly, 5 mm cylindrical liver cores were obtained with a surgical biopsy punch and sectioned to 200 μm slices using a Krumdieck tissue slicer (Alabama Research and Development, Munford, AL, USA) filled with carbonated Krebs-Henseleit buffer (pH 7.4, supplemented with 25 mM glucose). At this time point, some liver slices were snap-frozen in liquid nitrogen for RNA isolation. The rest was incubated in William’s E Medium (Gibco, Paisley, Scotland) supplemented with pen/strep in 6-well plates at 37°C/5% CO_2_/80% O_2_ under continuous shaking (70 rpm). 3 liver slices were incubated per well. After 1 h, medium was replaced with either fresh William’s E Medium 1% BSA in the presence or absence of a mix of 0.8 mM oleic acid and 0.02 mM BODIPY FL C12 (ThermoFisher Scientific, Breda, Netherlands) for imaging, or William’s E Medium 1% BSA in the presence or absence of a mixture of oleate and palmitate (ratio 2:1, total concentration 0.8 mM) for RNA and protein isolation. After overnight incubation, liver slices were snap-frozen in liquid nitrogen and stored in −80°C for RNA and protein isolation. Alternatively, liver slices were fixed for 1h in 3.7% formaldehyde, transferred into an 8-well removable chamber (ibidi, GmbH, Martinsried, Germany) and coated with vectashield. Slices were imaged on a Leica TCS SP8 X confocal. BODIPY FL C12 was excited at 488 nm and detected using HyD in a spectral window of 505-550 nm.

### Primary Hepatocytes

Buffers: Hanks:112 mM NaCl, 5.4 mM KCl, 0.9 mM KH_2_PO_4_, 0.7 mM Na_2_HPO_4_.12H_2_O. Hanks I: Hanks supplemented with 25 mM NaHCO_3_, 10 mM D-glucose, 0.5 mM EGTA at pH 7.42. Hanks II: same as Hanks I with the addition of 5 mM CaCl_2_. Krebs: 25 mM NaHCO_3_, 10 mM D-glucose, 10 mM Hepes. Hepatocyte culture medium: Williams E without phenol red (Fisher) supplemented with Primary hepatocyte maintenance supplement (Fisher). All buffers were saturated with carbogen before use.

Primary hepatocytes were prepared from *Hilpda*^Δhep^ and *Hilpda*^flox/flox^ mice. Briefly, mice were anesthetized with isoflurane. Livers were infused through the portal vein with Hanks buffer I for 10 min and with 100 mL of Hanks buffer II. Next, livers were infused with 200 mL Liver Digest Medium (Fisher). Livers were excised and washed in Krebs Buffer. Primary hepatocytes were passed through a 100 µm mesh and centrifuged at 450 rpm for 4 min at 4°C. Supernatant was discarded and cells were washed again in cold Krebs medium. Supernatant was discarded and cells were resuspended in hepatocyte culture medium with 5% FCS and seeded on collagen-coated 8-well µ-slide glass bottoms (Ibidi, Martinsried, Germany). After 2h medium was refreshed with hepatocyte culture medium with 10% FCS and left overnight. Next day, cells were treated with 20 µM Atglistatin (Sigma-Aldrich) or DMSO control and left overnight. Next morning, treatments were refreshed for 2h before adding a mixture of oleate and palmitate (ratio 2:1, total concentration 0.8 mM). Cells were lipid loaded for 6h and then fixed for 20 min in 3.7% paraformaldehyde. Lipid droplets were stained with 3 µg/mL BODIPY® 493/503 and mounted with vectashield for imaging. Cells were imaged on a Leica TCS SP8 X confocal.

BODIPY was excited at 488 nm and detected using HyD in a spectral window of 505-550 nm. Images were acquired 1024 × 1024 pixels with pinhole set at 1 airy unit (AU), pixel saturation was avoided. Images were processed and analyse with Fiji. Briefly, images were converted to binary images, watershed, and LD size and number was measured with particle analysis set 0.07 µm^2^-infinity

### TG quantification in HepG2 cells

HepG2 cells were seeded in 24-well plates. Next day, cells were transduced with Adenovirus-GFP (AV-*Gfp*) or Adenovirus-mHilpda (AV-*Hilpda*) at 5 × 10^6^ IFU/mL media in DMEM (Lonza, Verviers, Belgium) supplemented with 10% fetal calf serum (Lonza) and 1% penicillin/streptomycin (Lonza), from now on referred as complete DMEM, and left overnight. Recombinant adenoviruses were generated by cloning *GFP* or mouse *Hilpda* cDNA in human adenovirus type5 (dE1/E3). Expression regulated by CMV promoter. Viruses were produced and titrated by Vector Biolabs (Philadelphia, PA, USA). Cells were then incubated with DMEM 3% BSA and a mixture of oleate and palmitate (ratio 2:1, total concentration 1 mM) for 3 hours. Cells were washed twice with PBS and frozen in 25 mM Tris/HCl, 1mM EDTA, pH 7.5. TG quantification plates were thawed and a mixture of 4:1 tertiary butanol:methanol was added to cells, followed by an incubation of 10 minutes on a shaking platform. Plates were left to evaporate on a hot plate at 50°C. Next, 300 μL of triglyceride Liquicolor reagent (Human Diagnostics, Wiesbaden, Germany) was added to the cells and incubated for 10 minutes while shaking. 100 µL was transferred to a 96 well plate and absorption measured at 492 nm. A calibration curve of a standard solution was used to determine the TG content of the cells. The triglyceride content relative to protein content was then calculated. For protein quantification Pierce BCA kit (ThermoFisher Scientific) was used according to manufacturer’s protocol.

### LD count

HepG2 and Hepa1-6 cells were plated on collagen-coated 8-well µ-slide glass bottoms (ibidi, Martinsried, Germany). The next day, cells were transduced with AV-*Hilpda* in complete DMEM at 5×10^6^ IFU/mL media and left overnight. HepG2 cells were incubated with a mixture of oleate and palmitate (ratio 2:1, total concentration 0.8 mM) for 8h to promote LD formation. Hepa 1-6 cells were incubated with 1 mM oleate:palmitate for 24h. Cells were washed with PBS, fixed for 15 min with 3.7% formaldehyde, stained with 3 µg/mL BODIPY® 493/503 and Hoechst for 45 min, and mounted with Vectashield-H (Vector Laboratories). Cells were imaged on a Leica confocal TCS SP8 X system equipped with a 63× 1.20 NA water-immersion objective lens. BODIPY® 493/503 was excited at 488 nm and fluorescence emission was detected using internal Hybrid (HyD) in a spectral window of 505nm - 578nm. Images were acquired 1024 × 1024 pixels with pinhole set at 1 AU, pixel saturation was avoided. Images were processed and analyse with Fiji. Briefly, images were converted to binary, watershed and LD size and number was measured with particle analysis set 0.07 µm^2^-infinity.

### Western blot

Cell or tissue protein lysates were separated by SDS-PAGE on pre-cast 8-16% polyacrylamide gels and transferred onto nitrocellulose membranes using a Trans-Blot® Semi-Dry transfer cell (all purchased from Bio-Rad Laboratories), blocked in non-fat milk and incubated overnight at 4°C with primary antibody for HILPDA (1:750, Santa Cruz Biotechnology, sc-137518 or rabbit antisera against the C-terminal half (aa 37–64) of murine HILPDA generated by Pineda (Berlin, Germany) (23)), ACTIN (Cell Signaling Technology), ATGL (Santa Cruz Biotechnology), DGAT1 (Santa Cruz Biotechnology, sc-271934) or HSP90 (1:5000, Cell Signaling Technology, #4874). Membranes were incubated with a secondary antibody (Anti-rabbit IgG, HRP-linked Antibody, 7074, Cell Signaling Technology) and developed using Clarity ECL substrate (Bio-Rad Laboratories). Images were captured with the ChemiDoc MP system (Bio-Rad Laboratories).

### Plasmid constructs

Plasmids for *Plin2, Plin3, Gpat1, Gpat4, Dgat1, Dgat2* and *Hilpda* were constructed by fusing the full-length mouse cDNA into pEGFP-N2 (Clonetech, Mountain View, California, USA) and substituting the EGFP sequence by the sequence of the fluorescent proteins (FP) mCherry, sYFP2 or mEGFP. Briefly, RNA from mouse WAT or liver was reverse transcribed with First Strand cDNA synthesis kit (Thermo Scientific) and amplified with Phusion High fidelity DNA Polymerase (Thermo Scientific) using gene-specific primers. The PCR products were cloned into pEGFP-N2 vector using the XhoI and KpnI-HF or NheI and BamHI (New England Biolabs Inc.) restriction enzyme sites. Afterwards, MAX Efficiency ® DH5α(tm) Competent Cells (Invitrogen) were transformed by heat-shock and grown in Luria-Bertani (LB) agar plates with kanamycin (Sigma-Aldrich). The vector was isolated using Qiagen plasmid maxi kit (Qiagen) according to manufacturer instructions. The EGFP sequence was then excised from the pEGFP-N2 parent vector by enzyme digestion with KpnI-HF and NotI-HF. The vector was gel-purified with QIAquick Gel Extraction Kit (Qiagen) and the fragments of mCherry, sYFP2 or mEGFP were ligated into KpnI and NotI restriction enzyme site using T4 DNA ligase (Thermo Scientific). For plasmids of mGPAT1 and mGPAT4 the original pEGFP-N2 plasmid was used.

### Stimulated Emission Depletion (STED) microscopy

HepG2 cells were plated on collagen coated 8 well µ-slide glass bottom (ibidi, Martinsried, Germany). Next day cells were transfected with 750 ng of *Hilpda*_sYFP2 complexed to polyethylenimine (PEI) (Polyscience Inc., PA, USA) in serum free medium. After 6h, the transfection medium was changed to DMEM 1% FA-free BSA with 0.8 mM OA and 15 µM BODIPY C12 558/568. Cells were fixed after 18h lipid loading for 20 min in 3.7% PFA. Images were acquired on a Leica TCS SP8 STED microscope. A 100x 1.4 N.A. oil immersion objective was used in combination with a 5x optical zoom resulting in a pixel size of 23×23 nm. The pinhole was set at 0.9 AU and imaging speed at 700 Hz. For excitation of the fluorescent probes a white light laser line was used. HILPDA-sYFP2 and BODIPY-558 were excited at 470 nm and 558 nm respectively, and fluorescence emission was detected using HyD in a spectral window of 480-540 nm and 570-650 nm, respectively. The HILPDA-sYFP2 emission was partly depleted with the 592 depletion laser set at a 40% laser intensity with a power output of 1.3530 W. For both fluorophores the gating was set at 0.3-6.5 ns. Images were corrected for chromatic aberration and deconvolved using the Deconvolution Express modus in Huygens Professional Software (Scientific Volume Imaging B.V., Hilversum, the Netherlands).

### HILPDA and fluorescently labeled fatty acid colocalization

HepG2 cells were plated on collagen coated 8 well µ-slide glass bottom (Ibidi, Martinsried, Germany). Next day cells were transfected with Hilpda_mTurquoise2 plasmid complexed to PEI in serum-free DMEM. After 6 h, the medium was replaced by complete DMEM and left overnight. Cells were then incubated 16h with 0.6 mM oleate and 15 µM BODIPY C12 558/568 and next day for 20 min with QBT fatty acid uptake solution, which uses a BODIPY FL ®-dodecanoic acid fluorescent fatty acid analogue (BODIPY FL C12), prepared according to manufacturer’s protocol (Molecular Devices, California, USA). Cells were washed with PBS and fixed with 3.7% formaldehyde for 30 min, and mounted with vectashield (Molecular Devices, California, USA). Imaging was performed on a Leica TCS SP8 X system equipped with a 63x 1.20 NA water-immersion objective lens. Images were acquired sequentially 1024×1024 pixel scans with pinhole set at 1 AU. mTurquoise2 was excited at 440 nm and fluorescence emission was detected using internal HyD in a spectral window of 450-480 nm. BODIPY C12 558/568 was excited at 561 nm and fluorescence emission was detected using internal Hybrid (HyD) in a spectral window of 570-620 nm. BODIPY FL C12 was excited at 488 nm and fluorescence emission was detected using internal HyD in a spectral window of 505-558 nm. During image acquisition, fluorescence bleed-through and pixel saturation were avoided. All images were deconvolved using Deconvolution Express modus with Huygens Essential version 18.10 (<u>Scientific Volume Imaging</u>, The Netherlands). Further images were process with ImageJ. Briefly, channels were split, the entire cell was selected as a ROI, colocalization threshold was used to obtain colocalized pixels image and Mander’s and Pearson’s was measured using Coloc2 plugin.

### 2D Time Lapse

HepG2 cells were seeded on 15 µ-8 well glass bottom slide (Ibidi, Martinsried, Germany) and let grown overnight before transfection. Cells were transfected with 800 ng of mHilpda_mCherry plasmid complexed to polyethylenimine (PEI) (Polyscience Inc., PA, USA) in serum-free DMEM. After 6 h, the medium was replaced by complete medium. Next day cells were starved for 1h with HBSS 0.2% FA-free BSA. Medium was then replaced with QBT fatty acid uptake assay kit and after 4h incubation cells were imaged on a Leica TCS SP8 X system equipped with a 63x 1.20 NA water-immersion objective lens. Images were acquired sequentially using 512 × 512 pixels, and a total of 491 frames were acquired with a frame interval of ± 5 seconds. All images were deconvolved using Deconvolution Express modus with Huygens Essential version 18.10 (Scientific Volume Imaging, The Netherlands, http://svi.nl). Further images were processed with Fiji to assign different coloring LUTs for visualization.

### Lipophagy assay

HepG2 cells were transduced with AV-*GFP* or AV-*Hilpda* in complete DMEM at 5× 10^6^ IFU/mL media and left overnight. Next day, HepG2 cells were incubated with a mixture of 8 mM oleate:palmitate (ratio 2:1) for 8h to promote LD formation. 2h before collection, cells were treated with lysosomal inhibitor cocktail 20 mM ammonium chloride and 100 uM leupeptin. Cells were then lysed in RIPA buffer with protease and phosphatase inhibitors, and centrifuged at 10,000 rpm for 10 min.

### FRET-FLIM analysis

FRET is a process in which the excitation energy is transferred from a donor fluorophore to an acceptor chromophore in very close proximity (<10 nm). FRET determined using FLIM is independent of protein concentration, but very sensitive to the local microenvironment of the fluorophores. In FRET-FLIM, the fluorescence lifetime of the donor molecule is reduced in the presence of a nearby acceptor molecule, because energy transfer to the acceptor will introduce an additional relaxation path from the excited to the ground state of the donor (34).

HepG2 cells were cultivated in complete DMEM (at standard conditions (37 °C, 5% CO2, 95% humidified atmosphere). Cells were seeded on a rat tail collagen coated (Ibidi, Martinsried, Germany) 15 µ-8 well glass bottom slide (Ibidi, Martinsried, Germany) and let to grow for 24h before transfection. Transfections were performed with 800 ng of single or 1600 ng of mixed plasmid DNA complexed to polyethylenimine (PEI) (Polyscience Inc., PA, USA) in serum-free DMEM. After 5 h, the medium was replaced by serum free DMEM supplemented with 1% fatty acid free BSA (Roche Applied Sciences) and a mixture of oleate and palmitate (ratio 2:1, total concentration 0.8 mM) and left overnight. For imaging, medium was replaced with FluoroBrite DMEM supplemented with 1% BSA and 0.8 mM fatty acid mix.

Colocalization imaging was performed on a Leica TCS SP8 X system equipped with a 63x 1.20 NA water-immersion objective lens. Images were acquired sequentially 512× 512 pixels with pinhole set at 1 AU. mEGFP was excited at 488 nm and fluorescence emission was detected using internal HyD in a spectral window of 505-550 nm. mCherry was excited at 561 nm and detected using HyD in a spectral window of 580-650 nm. All images were deconvolved using Deconvolution Express modus with Huygens Essential version 18.10 (Scientific Volume Imaging, The Netherlands, http://svi.nl). Further images were process with Fiji. Briefly, brightness and contrast levels were adjusted and images were merged.

Förster resonance energy transfer-Fluorescence lifetime imaging microscopy (FRET-FLIM) was performed on a Leica TCS SP8 X confocal microscope. Donor and acceptor (mEGFP and mCherry, respectively) molecules were excited using a 40 MHz tunable supercontinuum laser at 488 nm and 561 nm, respectively. Fluorescence emission was detected using HyD detectors with 100 ps time resolution and collected in a spectral window of 505-550 nm for the donor (mEGFP) and 580-650 nm for the acceptor (mCherry). The signal output from the HyD detector was coupled to an external time-correlated single photon counting module (Becker&Hickl) for acquiring FLIM data. Typical images had 256 × 256 pixels (pixel size ± 300 nm), and the analogue to digital converter (ADC) was set to 256 time channels and FLIM images were acquired by imaging for 120 seconds per image. From the time resolved fluorescence intensity images, the fluorescence decay curves were calculated for each pixel and fitted with a double-exponential decay model using the SPCImage v7.1 software (Becker & Hickl). Fitting was performed without fixing any parameters. FRET-FLIM analysis provided fluorescence intensity as well as false-colored fluorescence lifetime images. The raw data was subjected to the following criteria to analyze and omit false positive negatives in the fluorescence lifetime scoring: minimum photon count per pixel of 1000 photons, 2 component analysis, goodness of fit (χ2<2) and fluorescence lifetime range of 500–3500 ps. For data analysis, we set pixel binning at 1 to have sufficient number of photons per pixel required for accurate fluorescence lifetime analysis.

### DGAT assay

Our protocol is a modification of the method described by McFie and Stone (26). HepG2 cells were seeded in 6-well plates at a density of 4 × 10^5^ cells/ well or in 60 × 15mm round cell culture dishes at a cell density of 3.5 × 10^6^ cells/dish in DMEM supplemented with 10% Fetal Calf Serum (FCS) and 1% Penicillin-Streptomycin (PS). Next day, cells were transduced with AV-*Gfp* or AV-*Hilpda* at 5×10^6^ PFU/mL medium. After 6h, the medium was changed to complete DMEM with 40 µM Atglistatin (3-(4′-(Dimethylamino)-[1,1′-biphenyl]-3-yl)-1,1-dimethylurea, Sigma-Aldrich) or control, and incubated overnight. Atglistatin is a specific high-affinity inhibitor of ATGL (25). For the samples treated with Atglistatin, Atglistatin was added again the next morning 2 h prior to cell lysate isolation. In addition, Atglistatin was added to the resuspension buffer during the fluorescence assay. Cells were detached with trypsin, washed, and resuspended in 100 μL of 50 mM Tris-HCl (pH 7.6)/250 mM sucrose buffer supplemented with protease inhibitors (Roche Diagnostics GmbH). Cells were disrupted by 20 passages through a 27-gauge needle. Prepared cell lysate samples were placed on a spinning wheel for 20 min at 4°C. Cell debris was pelleted by centrifugation at 2500 rpm for 5 min. The supernatant was transferred to a new tube and used for the assay. Protein concentration was determined using a Pierce BCA kit (Thermo Fisher Scientific). A master mix containing 20 μL of 1 M Tris-HCl (pH 7.6), 4 μL of 1 M MgCl2, 10 μL of 4 mM DOG (Sigma-Aldrich), 10 μL of 12.5 mg/mL BSA, 10 μL of 500 μM NBD-palmitoyl CoA (Avanti Polar Lipids), and 96 μL of water per reaction, was prepared. Volumes were scaled up proportionally to accommodate the desired number of reactions. The master mix was protected from direct light during the entire experiment by wrapping the glass test tubes in aluminium foil. Assays were performed in 13 × 100 mm glass KIMAX Test Tubes with Teflon Liner Caps (DWK Life Sciences, Kimble) in a final reaction volume of 250 μL. A master mix volume of 150 μL was aliquot per test tube, and tubes pre-incubated in a 37°C water bath for 2 min. The reaction was started by adding 300 ug in 100 μL of protein sample and incubated at 37°C Shaking Water Bath (GFL Gesellschaft für Labortechnik mbH, Product No. 1086) for 30, 90 and/or 180 min with steady shaking at 60 rpm. For the co-treatment with Atglistatin (40 μM), DGAT1 inhibitor (A922500, 1 μM) and DGAT2 inhibitor (PF-06424439, 40 μM), the incubation was carried on for 180 min. The reaction was terminated by adding 4 mL CHCl_3_/methanol (2:1, v/v) and 800 μL of water mixed by vortex. After 1 h, the test tubes containing samples were re-vortexed and centrifuged at 3,000 rpm for 5 min to separate aqueous and organic phases. The upper aqueous phase was aspirated, and the organic phase dried under stream of nitrogen. To help the solvents evaporate faster, the test tubes were placed in a thermal block pre-warmed to 54°C. Lipids were finally resuspended in 50 μL CHCl3/methanol (2:1) and stored at −20°C overnight. Samples were vortexed and re-centrifuged at 3,000 rpm for 2 min, before being spot on channelled 20 × 20 cm TLC plates with pre-adsorbent silica gel HLF zone (Analtech). The TLC plates were developed in the solvent system containing hexane/ethyl ether/acetic acid (80:20:1, v/v/v). The plates were air dried for 1 h before quantification of reaction products.

The newly synthesized NBD-TG was analysed with a ChemiDocTM MP molecular imaging system (Bio-Rad Laboratories, Inc.), and fluorescence was quantified with Quantity One software 4.1 (Bio-Rad Laboratories, Inc.). The excitation and emission wavelengths of NBD are 465 nm and 535 nm, respectively. Extinction source UV Trans illumination and Standard Emission Filter, together with Application SYBER Green and Applied (UV Trans Orange) Flat Field, were used. Data is presented as arbitrary fluorescence intensity units.

### Microarray analysis

Microarray analysis was performed on Hepa1-6 hepatoma cells incubated with different fatty acids. RNA was purified with RNeasy Minikit columns (Qiagen) and analysed for quality with RNA 6000 Nano chips on the Agilent 2100 bioanalyzer (Agilent Technologies, Amsterdam, The Netherlands). One microgram of RNA was used for cDNA synthesis using the First Strand cDNA synthesis kit (Thermo Scientific). Purified RNA (100 ng) was labeled with the Ambion WT expression kit (Invitrogen) and hybridized to an Affymetrix Mouse Gene 1.1 ST array plate (Affymetrix, Santa Clara, CA). Hybridization, washing, and scanning were carried out on an Affymetrix GeneTitan platform. Scans of the Affymetrix arrays were processed using packages from the Bioconductor project. Arrays were normalized using the robust multi-array average method (4, 22). Probe sets were defined by assigning probes to unique gene identifiers, e.g., Entrez ID (10). The total gene set (24,973 probe sets) was filtered to only include genes with mean signal > 20, yielding 10,379 genes. Microarray data were submitted to the Gene Expression Omnibus (accession number pending).

### Statistical analysis

Details of statistical analyses are given in the figure legends. Statistical analyses were carried out using an unpaired Student’s t test or two-way ANOVA. A value of p<0.05 was considered statistically significant.

## RESULTS

### Hilpda expression is induced by fatty acids in hepatoma cells

Previously, we and others found that expression of *Hilpda* is induced by fatty acids in cancer cells and macrophages (23, 35, 36). To examine if *Hilpda* expression is regulated by fatty acids in hepatoma cells, we treated mouse Hepa1-6 cells for 6h with different types of fatty acids: cis-unsaturated (oleate), trans-unsaturated (elaidate), or saturated (palmitate). Besides *Plin2, Hilpda* was one of the 65 genes that was induced at least 1.5 fold by palmitate (3.3 fold), oleate (1.5 fold) and elaidate (3.9 fold) (Figure 1a). The regulation of *Hilpda* strongly resembled the pattern observed for *Plin2* (Figure 1b). To further investigate the induction of *Hilpda* by fatty acids, different hepatoma cell types were treated with a mixture of oleate and palmitate. In mouse Hepa1-6, rat FAO, and human HepG2 cells, a mixture of oleate and palmitate significantly induced *Hilpda* mRNA, along with *Plin2* (Figure 1c). In Hepa1-6 cells, the mixture of oleate and palmitate also increased HILPDA protein. (Figure 1d). These data indicate that *Hilpda* expression is induced by fatty acids in liver cells.

**Figure 1:**
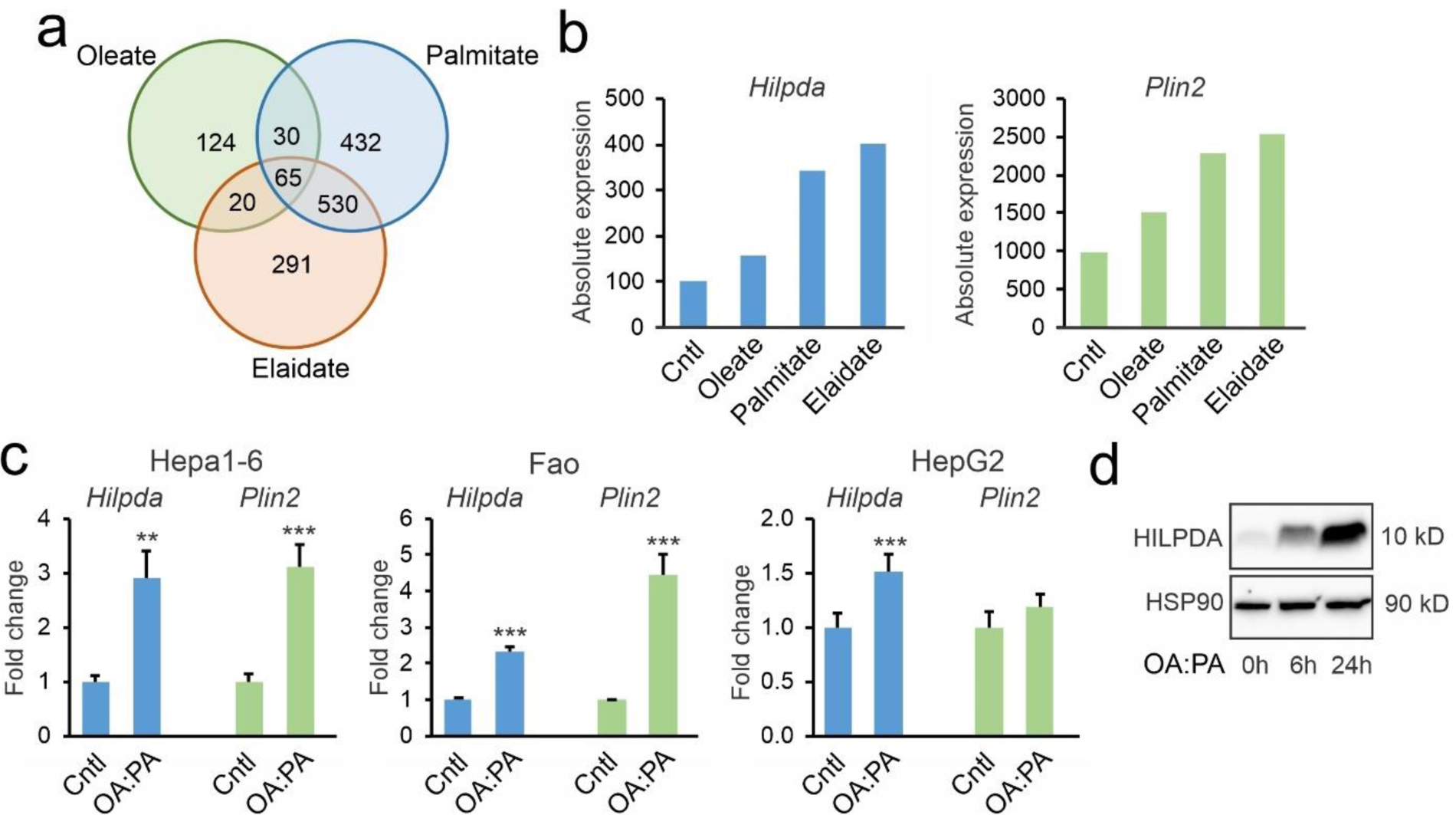
*Hilpda* expression is induced by fatty acids in various hepatoma cell lines. a) Venn diagram of upregulated genes (fold change>1.5) in murine Hepa1-6 hepatoma cells treated with different fatty acids (500 µM) for 6h. b) Relative changes in *Hilpda* and *Plin2* mRNA in Hepa1-6 cells treated with different fatty acids (500 µM) for 6h. c) Relative changes in *Hilpda* and *Plin2* mRNA in Hepa1-6, Fao and HepG2 hepatoma cells treated for 24h with a 2:1 mixture of oleate and palmitate (total concentration 1.2 mM). d) HILPDA protein levels in Hepa1-6 cells treated with a 2:1 mixture of oleate and palmitate for different duration. Bar graphs are presented as mean ±SD. Asterisk indicates significantly different from control-treated cells according to Student’s t test; **P < 0.01; ***P < 0.001.

### HILPDA deficiency modestly decreases liver triglyceride storage in mice with NASH

Previously, it was found that hepatocyte-specific deficiency of HILPDA reduced hepatic triglyceride levels under chow-fed conditions, although not after chronic high fat feeding (12). A possible reason for the inconsistent effect of HILPDA deficiency on liver triglycerides is the relatively low *Hilpda* expression in liver. To identify conditions where deficiency of HILPDA may be expected to have a larger effect, we screened mouse liver transcriptome data for upregulation of *Hilpda*. Interestingly, hepatic *Hilpda* mRNA levels were increased during non-alcoholic steatohepatitis (NASH) caused by feeding mice a methionine and choline deficient diet (Figure 2a, based on GSE35961). Accordingly, we hypothesized that the effect of HILPDA deficiency may be more pronounced during NASH. Hepatocyte-specific HILPDA-deficient mice were generated by crossing *Hilpda*^flox/flox^ mice with mice expressing Cre-recombinase driven by the albumin promoter. To induce NASH, *Hilpda*^Δhep^ and *Hilpda*^flox/flox^ mice were fed a high-fat diet deficient in methionine and choline for 11 weeks (HFmcd), using a low-fat (LF) diet as control. After 11 weeks, hepatic expression of *Hilpda* was significantly higher in mice fed the HFmcd than the LF diet, and was significantly lower in *Hilpda*^Δhep^ than in *Hilpda*^flox/flox^ mice, which was accompanied by a compensatory increase in *G0s2* mRNA (Figure 2b). HILPDA protein levels were also markedly reduced in livers of *Hilpda*^Δhep^ compared to *Hilpda*^flox/flox^ mice (Figure 2c). Mice fed HFmcd were significantly lighter than the mice fed LFD, but no differences were observed between the genotypes (Figure 2d). Similarly, weight of the epididymal fat pad was significantly lower in the mice fed HFmcd, but no differences were observed between *Hilpda*^Δhep^ and *Hilpda*^flox/flox^ mice (Figure 2e). Intriguingly, in mice fed HFmcd but not the LF diet, the weight of the liver was modestly but significantly lower in *Hilpda*^Δhep^ than in *Hilpda*^flox/flox^ mice (Figure 2f). Consistent with a stimulatory effect of HILPDA on liver fat, hepatic triglyceride levels were modestly but significantly lower in *Hilpda*^Δhep^ compared to *Hilpda*^flox/flox^ mice, both on the HFmcd and LF diet (Figure 2g).

**Figure 2:**
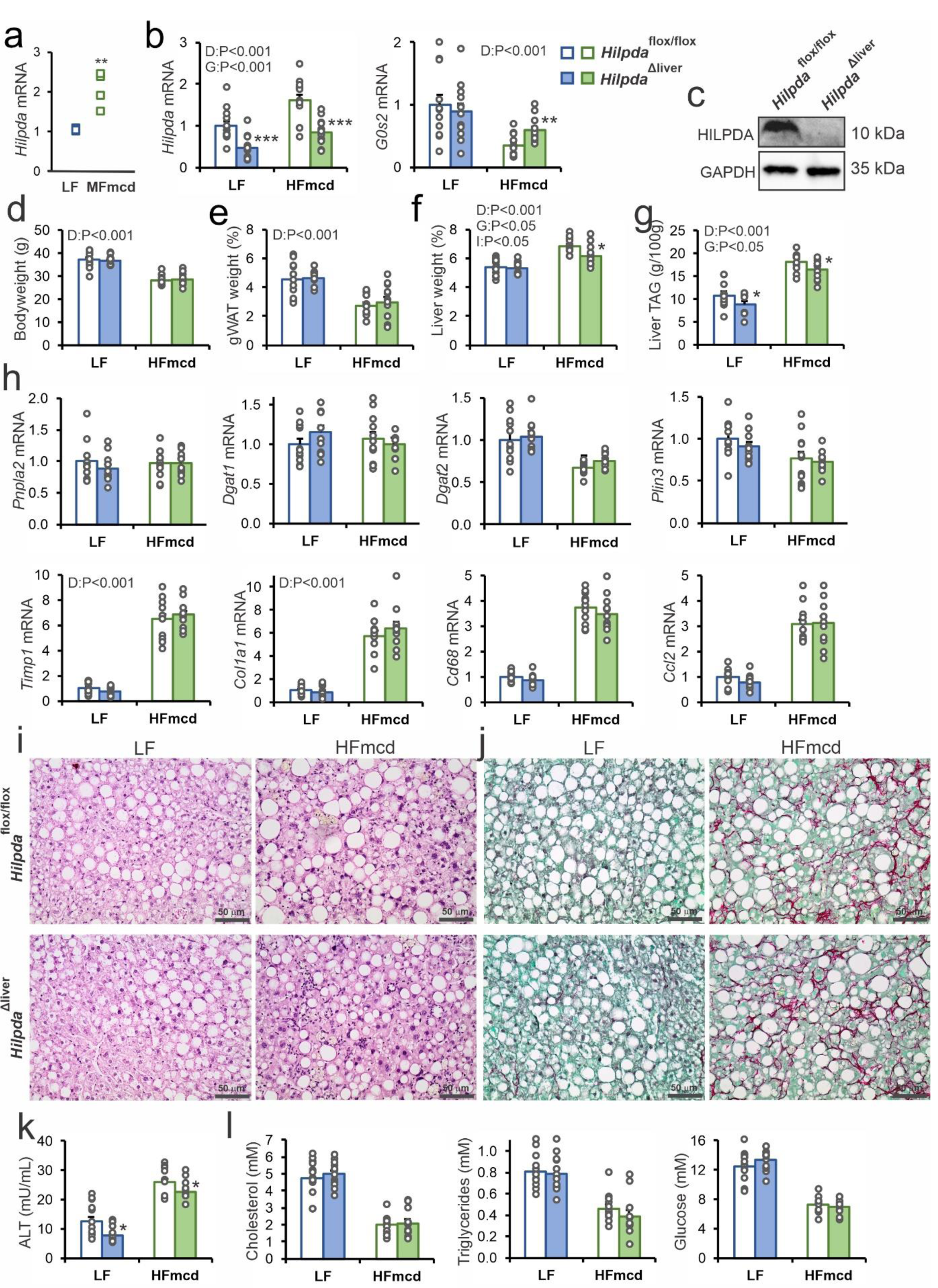
Effect of hepatocyte-specific HILPDA deficiency in mice with NAFLD. a) Upregulation of hepatic *Hilpda* mRNA by methione and choline-deficient diet (GSE35961). b) *Hilpda* and *G0s2* mRNA levels in livers of *Hilpda*^Δhep^ and *Hilpda*^flox/flox^ mice fed a low fat diet (LF) or high fat diet deficient in methionine and choline (HFmcd). c) HILPDA protein levels in livers of *Hilpda*^Δhep^ and *Hilpda*^flox/flox^ mice fed a low fat diet. d) Body weight. e) Gonadal adipose tissue weight. f) Liver weight. g) Liver triglyceride levels. h) mRNA levels of various LD-associated proteins, inflammatory markers. and fibrosis markers. i) H&E staining. j) Syrius Red staining. k) Plasma ALT levels. l) Plasma levels of various metabolites. Data are mean ± SEM; N=12 mice/group. Asterisk indicates significantly different from *Hilpda*^flox/flox^ mice according to Student’s t test; *P < 0.05; **P<0.01; ***P < 0.001.

HILPDA deficiency did not have any significant effect on mRNA levels of *Pnpla2, Dgat1, Dgat2*, and *Plin3* (Figure 2h). Also, despite a marked induction by HFmcd of the expression of macrophage/inflammatory markers *Cd68* and *Ccl2*, and fibrosis markers *Timp1* and *Col1a1*, no significant differences were observed between *Hilpda*^Δhep^ and *Hilpda*^flox/flox^ mice (Figure 2h). Histological analysis by H&E (Figure 2i) and Syrius Red (Figure 2j) staining indicated that mice fed HFmcd exhibited classical features of NASH, including ballooning, inflammation, steatosis, and fibrosis. However, no clear and consistent differences were visible between *Hilpda*^Δhep^ and *Hilpda*^flox/flox^ mice. By contrast, and in agreement with the liver triglyceride levels, plasma ALT levels were modestly but significantly lower in *Hilpda*^Δhep^ compared to *Hilpda*^flox/flox^ mice, both on the HFmcd and LF diet (Figure 2k). Finally, no significant differences in plasma cholesterol, triglycerides and glucose were observed between *Hilpda*^Δhep^ and *Hilpda*^flox/flox^ mice on either diet (Figure 2l). Overall, these data indicate that hepatocyte-specific HILDPA deficiency causes a modest but significant decrease in hepatic triglyceride storage, liver weight, and plasma ALT levels, without having a clear impact on features of NASH and various metabolic parameters.

### HILPDA promotes lipid storage at least in part independently of ATGL

To further study the functional role of HILPDA in liver cells, we used precision cut liver slices and primary hepatocytes. These model systems were chosen because they both express very high levels of *Hilpda* compared to mouse liver (Figure 3a). Liver slices were prepared from *Hilpda*^Δhep^ and *Hilpda*^flox/flox^ mice and incubated overnight with a mixture of oleate and palmitate, as well as with BODIPY FL C12, followed by visualization of the stored fatty acids using fluorescence confocal microscopy. Levels of *Hilpda* mRNA were about 60% lower in *Hilpda*^Δhep^ than *Hilpda*^flox/flox^ liver slices, and HILPDA protein levels were also markedly reduced (Figure 3b). Consistent with a stimulatory effect of HILPDA on lipid storage, hepatocyte-specific deficiency of *Hilpda* led to a marked reduction in BODIPY FL accumulation in LD (Figure 3c).

**Figure 3:**
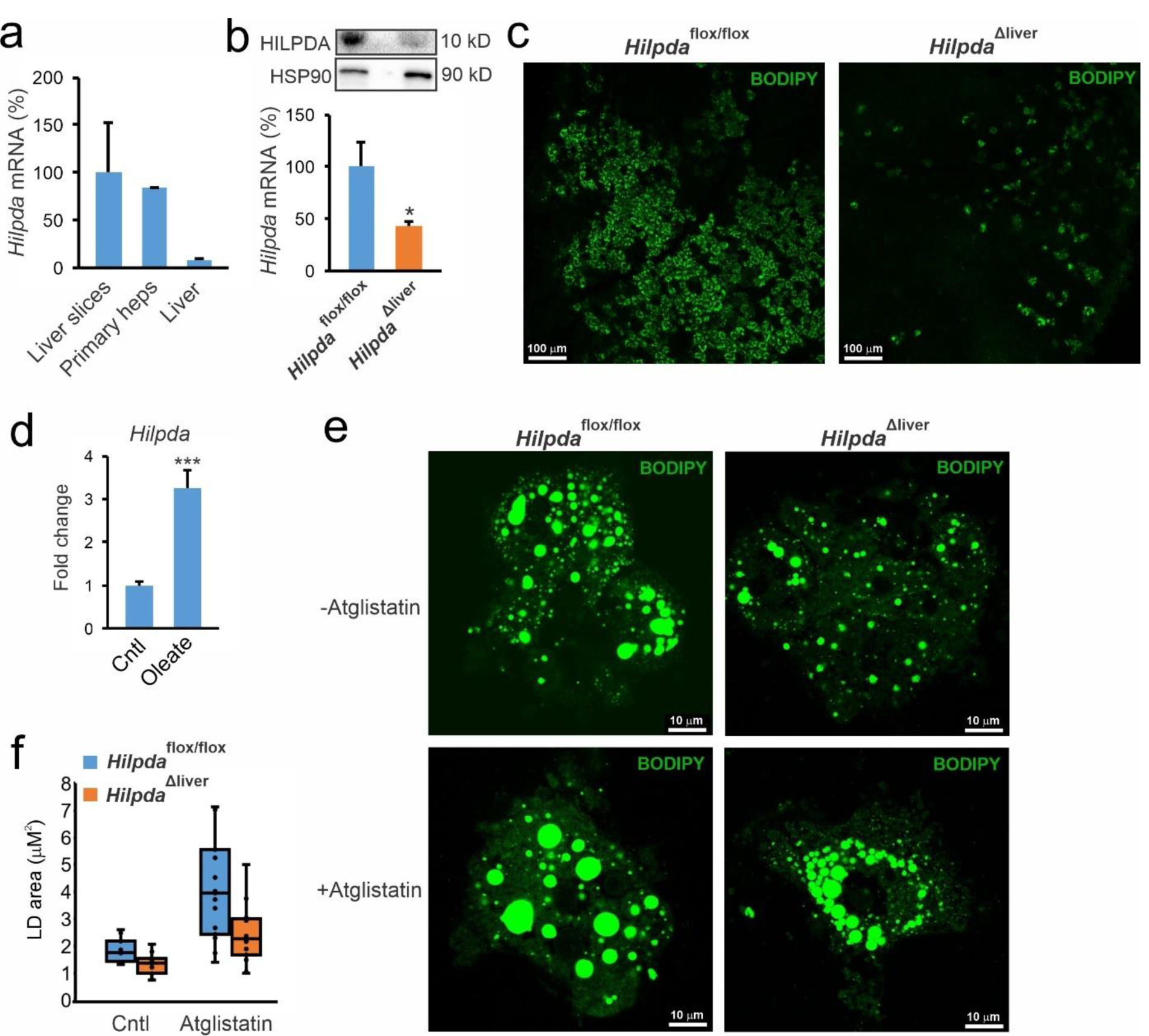
HILPDA stimulates lipid droplet formation partly independently of ATGL. a) Relative *Hilpda* mRNA levels in mouse precision cut liver slices, mouse primary hepactocytes, and mouse liver. b) HILPDA protein levels (top panel) and relative *Hilpda* mRNA levels (lower panel) in liver slices prepared from *Hilpda*^Δhep^ and *Hilpda*^flox/flox^ mice. Bar graphs are presented as mean ±SD. Asterisk indicates significantly different from *Hilpda*^flox/flox^ mice according to Student’s t test; *P < 0.05. c) Confocal microscopy of liver slices prepared from *Hilpda*^Δhep^ and *Hilpda*^flox/flox^ mice and incubated overnight with 800 µM oleate and 20 µM BODIPY FL C12. λ_ex_: 488nm, λ_em_: 550-595 nm. d) *Hilpda* mRNA expression in wildtype mouse primary hepatocytes treated with oleate (500 µM) for 24h. e) BODIPY 493/503 staining of primary hepatocytes prepared from *Hilpda*^Δhep^ and *Hilpda*^flox/flox^ mice and incubated overnight with 0.8 mM oleate:palmitate mix (2:1) acid in the presence or absence of Atglistatin (20 µM). λ_ex_: 488nm, λ_em_: 550-595 nm. f) Quantification of the lipid droplet area. The total number of lipid droplets analyzed per condition varied between 1017 and 1958. Two-way ANOVA revealed significant effects for Atglistatin (P<0.001) and genotype (P<0.001), but not for an interaction effect.

We next moved to primary hepatocytes. In these cells, *Hilpda* mRNA was significantly induced by fatty acids (Figure 3d). To examine the effect of HILPDA deficiency on lipid storage, primary hepatocytes of *Hilpda*^Δhep^ and *Hilpda*^flox/flox^ mice were incubated overnight with a mixture of oleate and palmitate, followed by visualization of lipid storage by BODIPY 493/503 staining and fluorescence confocal microscopy. Again, consistent with a stimulatory effect of HILPDA on lipid storage, LD were considerably smaller in *Hilpda*^Δhep^ than *Hilpda*^flox/flox^ primary hepatocytes (Figure 3e). Quantification of the images revealed a significantly lower LD area in the *Hilpda*^Δhep^ than *Hilpda*^flox/flox^ hepatocytes (Figure 3f). Previously, we and others found that HILPDA inhibits ATGL (28, 44). To investigate if the effect of loss of HILPDA on lipid storage in hepatocytes is mediated by hyperactivation of ATGL, we cotreated *Hilpda*^Δhep^ and *Hilpda*^flox/flox^ hepatocytes with the ATGL inhibitor Atglistatin. While Atglistatin and *Hilpda* genotype significantly increased the LD area, no statistical interaction was observed between Atglistatin treatment and *Hilpda* genotype (Figure 3e,f), suggesting no functional interaction between ATGL and HILPDA. These data suggest that HILPDA promotes lipid storage at least partly via an ATGL-independent mechanism.

### HILPDA preferentially associates with newly synthesised lipid droplets and active lipid droplets

To better understand the mechanism by which HILPDA promotes lipid storage in liver cells, we first investigated HILPDA localization. Previously, HILPDA protein was shown to partly localize to LD and to the LD-ER interface (11, 12, 15, 23). Cellular fractionation confirmed the association of HILPDA with lipid droplet in mouse liver, which intriguingly was reduced by fasting (Supplemental figure 1a). The fractionation of the liver was confirmed by immunoblot of marker genes, showing the pronounced activation of autophagy by fasting in the LD fraction (Supplemental figure 1b).

To better zoom in on the intracellular localization of HILPDA, we used stimulated emission depletion (STED) microscopy, which generates images of very high spatial resolution (± 70 nm). HepG2 cells were transfected with HILPDA fused to sYFP2 and the LD were visualized by lipid loading the cells with a mix of oleate and BODIPY C12 558/568. Interestingly, while some HILPDA was observed around LD, most of the HILPDA was localized in the perinuclear area, presumably representing the ER, where triglyceride synthesis occurs (Figure 4a). Interestingly, many LD were not surrounded by HILPDA. Visualization of the ER via co-transfection with pDsRed2-ER verified the partial localization of HILPDA in the ER (Figure 4b).

**Figure 4:**
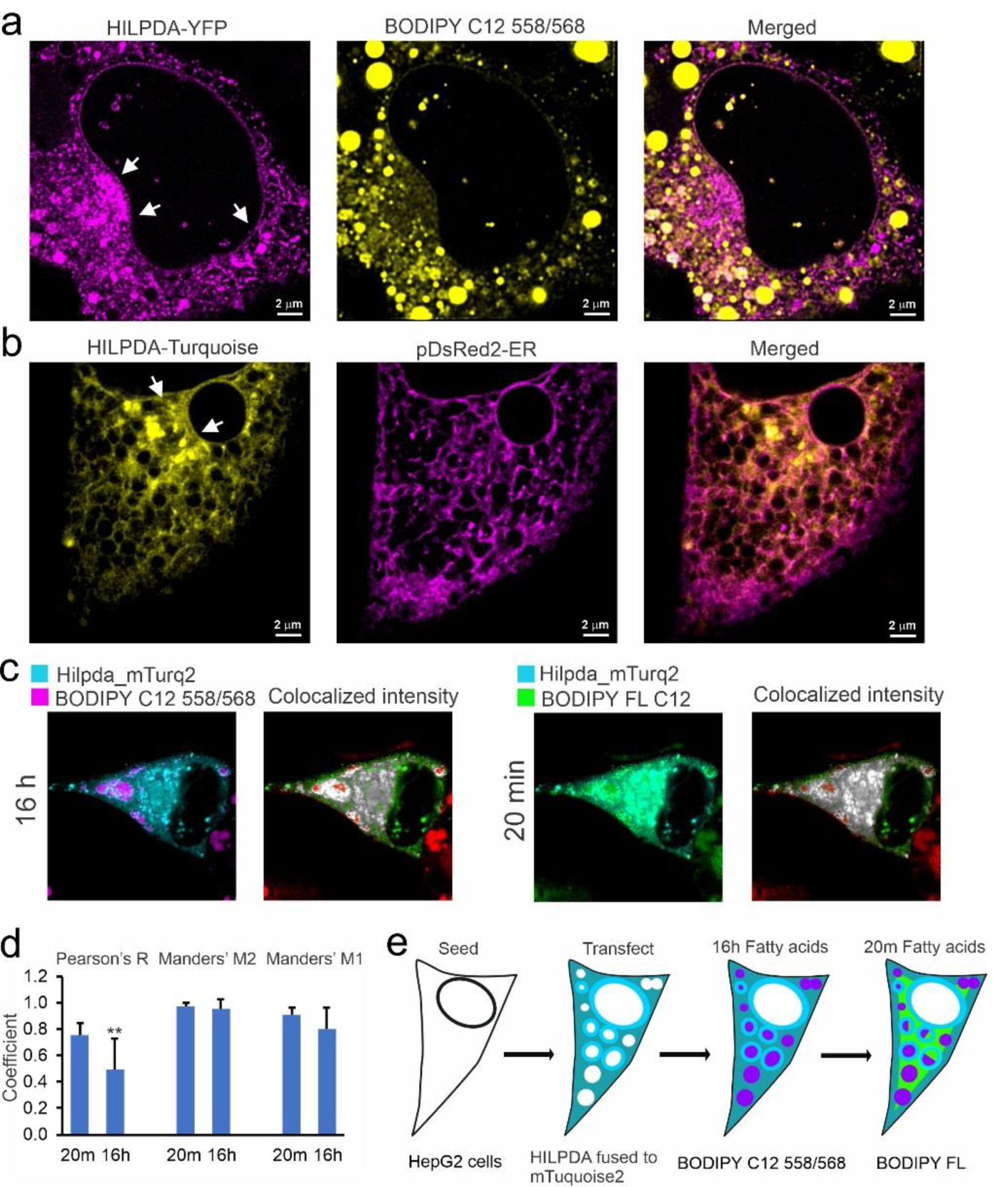
HILPDA is primarily localized to the perinuclear area and preferentially associated with new fat. a) STED microscopy of HepG2 cells transfected with HILPDA fused to sYFP2 and lipid loaded with 0.8 mM oleate and 15 µM BODIPY C12 558/568 for 18h. λ_ex_: 470 nm (YFP) and 558 nm (BODIPY-558/568). λ_em_: 480-540 nm (sYFP2) and 570-650 nm (BODIPY 558/568). Left panel: HILPDA-sYFP2; middle panel: BODIPY-558, right panel: overlay. Arrows indicate perinuclear area. b) HepG2 cells cotransfected with HILPDA fused to Turquoise2 and with ER marker pDsRed2-ER (ClonTech) followed by incubation with 0.8mM oleate overnight.Left panel: HILPDA-Turquoise2; middle panel: ER marker pDsRed2-ER, right panel: overlay. Arrows indicate perinuclear area. c) Confocal microscopy of HepG2 cells transfected with HILPDA fused to Turquoise2 and lipid loaded with 0.6 mM oleate and 15 µM BODIPY C12 558/568 for 16h, followed by incubation for 20 min with BODIPY FL C12 and fixed with 3.7% PFA. λ_ex_: 440 nm (mTurquoise2), 561 nm (BODIPY 558/568), and 488 nm (BODIPY FL). λ_em_: 450-480 nm (mTurquoise2), 570-620 nm (BODIPY 558/568), and 505-558 nm (BODIPY FL). Colocalized pixels of HILPDA and Fluorescent fatty acids are represented on gray scale, higher colocalization is depicted with lighter pixels; non-colocalized HILPDA pixels are coloured green; whereas non-colocalized fluorescent fatty acid pixels are coloured red. d) Mean Pearson’s correlation coefficient R, and mean Manders’ colocalization coefficients M2 and M1. Asterisk indicates significantly different according to Student’s t test; P<0.01. e) Schematic depiction of the set-up and outcomes of the above experiments.

To examine the dynamics of the association of HILPDA with LD, we performed time-lapse fluorescence imaging in HepG2 cells transfected with HILPDA-sYFP2 and incubated with BODIPY FL C12 (Supplemental Video 1). Intriguingly, HILPDA was mainly present around LD that are being lipolyzed (disappear) and remodelled (form new LD). Little to no HILPDA was observed around stable LD. These data suggest a role of HILPDA in LD remodelling in liver cells.

To better characterize the functional role of HILPDA in LD homeostasis in hepatocytes, we transfected HepG2 cells with HILPDA fused to mTurquoise2 and treated the cells with two labelled fatty acids that could be visualized separately using different channels. One labelled fatty acid (BODIPY C12 558/568) was added for 16 hours, while the other labelled fatty acid (BODIPY FL C12) was added for 20 min, after which cells were fixed (Figure 4c). Colocalization was evaluated by Manders Colocalization Coefficients and Pearson Correlation Coefficient (Figure 4d). Manders Colocalization Coefficients are measurements of co-occurrence, which is the spatial overlap of two probes. Pearson Correlation Coefficient is a measurement of correlation, which evaluates the spatial overlap and signal proportionality. Analysis of the confocal images showed that HILPDA colocalizes almost entirely with the old and newly added fatty acids (M2: 96-97% respectively), and that the proportion of fatty acids that colocalized with HILPDA is greater for the newly added fatty acids than for the fatty acids added the day before (new M1:91% vs. old M1:80%). In line with this, the Pearson Correlation Coefficient was significantly higher for the newly added fatty acids than for the fatty acids added the day before (Figure 4d). These data indicate that HILPDA more strongly correlates with newly added fatty acids, which in turn suggests that HILPDA preferentially colocalizes with newly synthesized triglycerides. A schematic depiction of the set-up and outcomes of the above experiments is presented in Figure 4e.

### HILPDA increases DGAT activity and DGAT1 levels

To further study how HILPDA promotes lipid storage, we overexpressed HILPDA in HepG2 cells via transduction with an adenoviral vector expressing *Hilpda*. HILPDA overexpression effectively raised HILPDA protein levels (Figure 5a), and was associated with a significantly increase in triglyceride levels (Figure 5b). BODIPY staining confirmed a significant increase in LD in HepG2 transduced with AV-*Hilpda* (Figure 5c). Quantification analysis showed that AV-*Hilpda* significantly increased the volume of the LD in both HepG2 and Hepa1-6 cells (Figure 5d). These data indicate that HILPDA overexpression promotes lipid storage in HepG2 cells.

**Figure 5:**
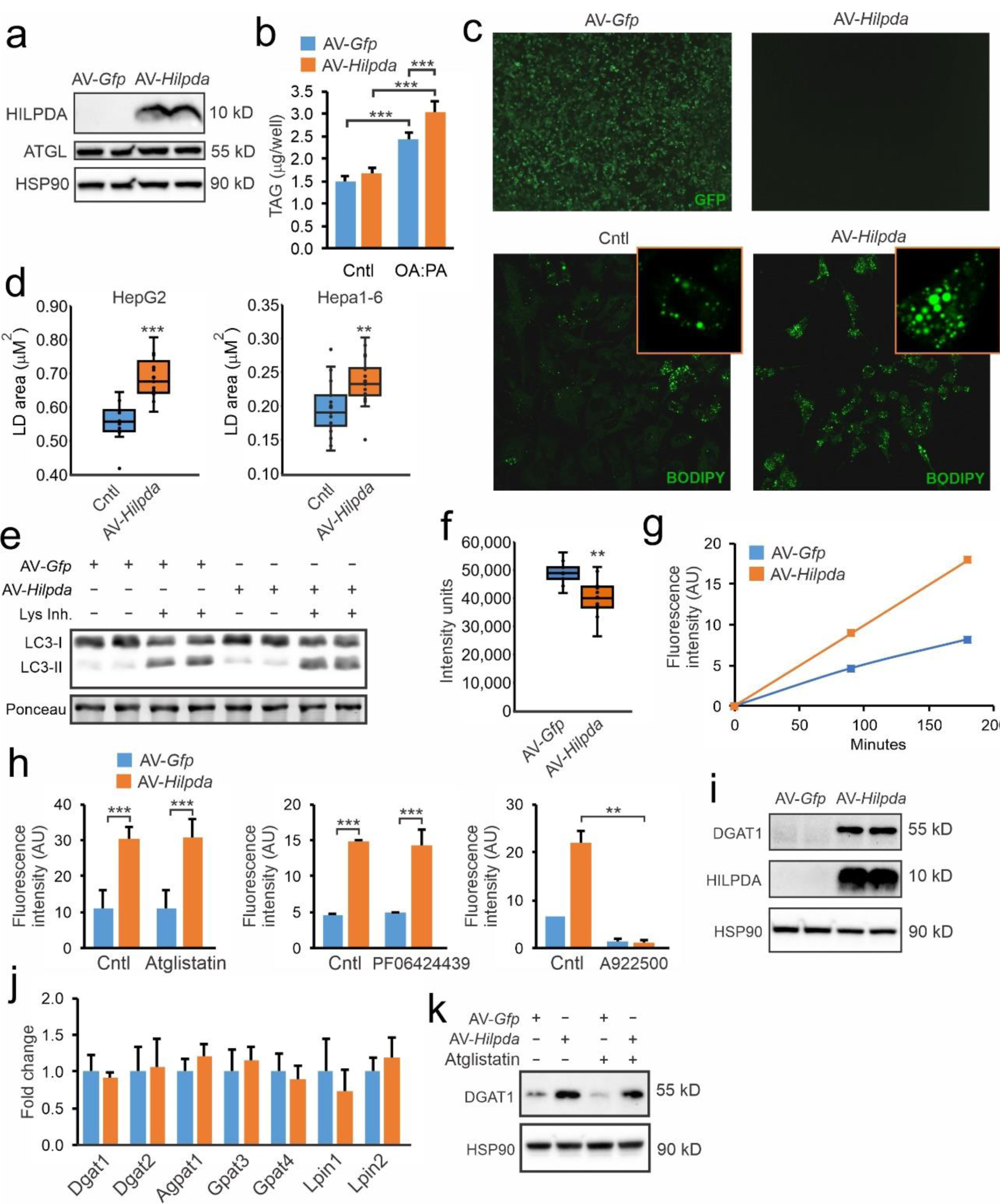
HILPDA overexpression promotes LD storage and increases DGAT1 levels in HepG2 cells. HepG2 cells were transduced with AV-*Hilpda*, AV-*GFP*, or non-transduced and treated with oleate:palmitate (2:1 ratio). a) HILPDA protein levels. b) Triglyceride content in HepG2 cells incubated with serum free DMEM or 3h in 1 mM oleate:palmitate. c) GFP fluorescence and BODIPY 493/503 staining. d) Quantification of LD size in HepG2 treated with 0.8mM oleate:palmitate for 8h and Hepa 1-6 cells treated with 1mM for 24h. e) LC3-I and LC3-II protein levels in HepG2 cells lipid loaded with 0.8mM oleate:palmitate for 8h, in the presence and absence of lysosomal inhibitors cocktail. f) Total DAG levels as determined by lipidomics in HepG2 cells incubated with 0.8mM oleate:palmitate for 5h. g) Time course of DGAT activity in HepG2 cells. h) DGAT activity in HepG2 cells in the presence and absence of ATGL, DGAT2 and DGAT1 inhibitor. i) DGAT1 and HILPDA protein levels in HepG2 cells. j) mRNA levels of selected genes. k) DGAT1 protein levels in HepG2 cells in presence and absence of Atglistatin.

Based on the finding that HILPDA increases lipid accumulation partly independent of ATGL, we considered the possibility that HILPDA may target the lipophagy pathway (31). However, accumulation of the autophagosome marker LC3-II in presence of lysosomal inhibitors was comparable between control and AV-*Hilpda* cells indicating that increased LD content in AV-*Hilpda* cells is not due to stimulation of lipophagy (Figure 5e). Alternatively, we considered that HILPDA may promote the synthesis and/or storage of triglycerides. Interestingly, lipidomics indicated that AV-*Hilpda* significantly decreased levels of diacylglycerols (Figure 5f), but did not significantly affect levels of other major lipid species (data not shown). Accordingly, we hypothesized that HILPDA might stimulate the activity of diacylglycerol acyltransferase (DGAT), which catalyzes the last and purportedly the rate-limiting step in the formation of triglycerides, using diacylglycerol and acyl-CoA as substrates. To determine a possible stimulatory effect of HILPDA on DGAT activity, we measured the synthesis of fluorescently labelled triglycerides from fluorescent NBD-palmitoyl-CoA and 1,2 dioleoyl-sn-glycerol in lysates of HepG2 cells transduced with AV-*Hilpda* or AV-*Gfp*. Strikingly, the DGAT-mediated incorporation of fluorescent NBD-palmitoyl-CoA into triglycerides, as determined by quantification of TLC plates, was markedly increased by HILPDA overexpression (Figure 5g). This increase in triglyceride synthesis in HepG2 cells was unaltered in the presence of Atglistatin, suggesting it is independent of ATGL (Figure 5h). DGAT activity is catalyzed by two different isozymes: DGAT1 and DGAT2 (3). Whereas DGAT2 has a preference for endogenously synthesized fatty acids, DGAT1 mainly esterifies exogenous fatty acids to diacylglycerol (30, 37, 38). Strikingly, the increase in triglyceride synthesis by HILPDA overexpression was unaltered by the DGAT2 inhibitor PF-06424439 but completely suppressed by the DGAT1 inhibitor A922500. Interestingly, the induction of DGAT activity in HepG2 cells by HILPDA overexpression was accompanied by a significant increase in DGAT1 protein levels (Figure 5i). The increase in DGAT1 protein was not associated with any change in mRNA levels of DGAT1, DGAT2 or other relevant proteins (Figure 5j), and was independent of ATGL activity (Figure 5k). The specificity of the DGAT1 antibody is shown in supplemental figure 2a. Neither the DGAT1 inhibitor nor the DGAT2 inhibitor affected endogenous HILPDA levels, as tested in Hepa1-6 cells (Supplemental figure 2b and 2c).

To assess the effect of HILPDA overexpression on hepatic lipid levels and DGAT1 protein levels in vivo, we used samples from a previous study in which mice infected mice with AAV expressing HILPDA(24). AAV-mediated HILPDA overexpression markedly changed the lipidomic profile in liver (Figure 6a) and markedly increased hepatic triglyceride levels (Figure 6b). In fact, out of 1480 different lipids, the 30 most significantly altered lipids by HILPDA overexpression were nearly all triglycerides, the remainder being cholesteryl-esters (Figure 6c). Consistent with the data in HepG2 cells, HILPDA overexpression increased DGAT1 protein levels in mouse liver (Figure 6d). These data suggest that HILPDA promotes triglyceride storage in liver concurrent with an increase in DGAT1 protein.

**Figure 6:**
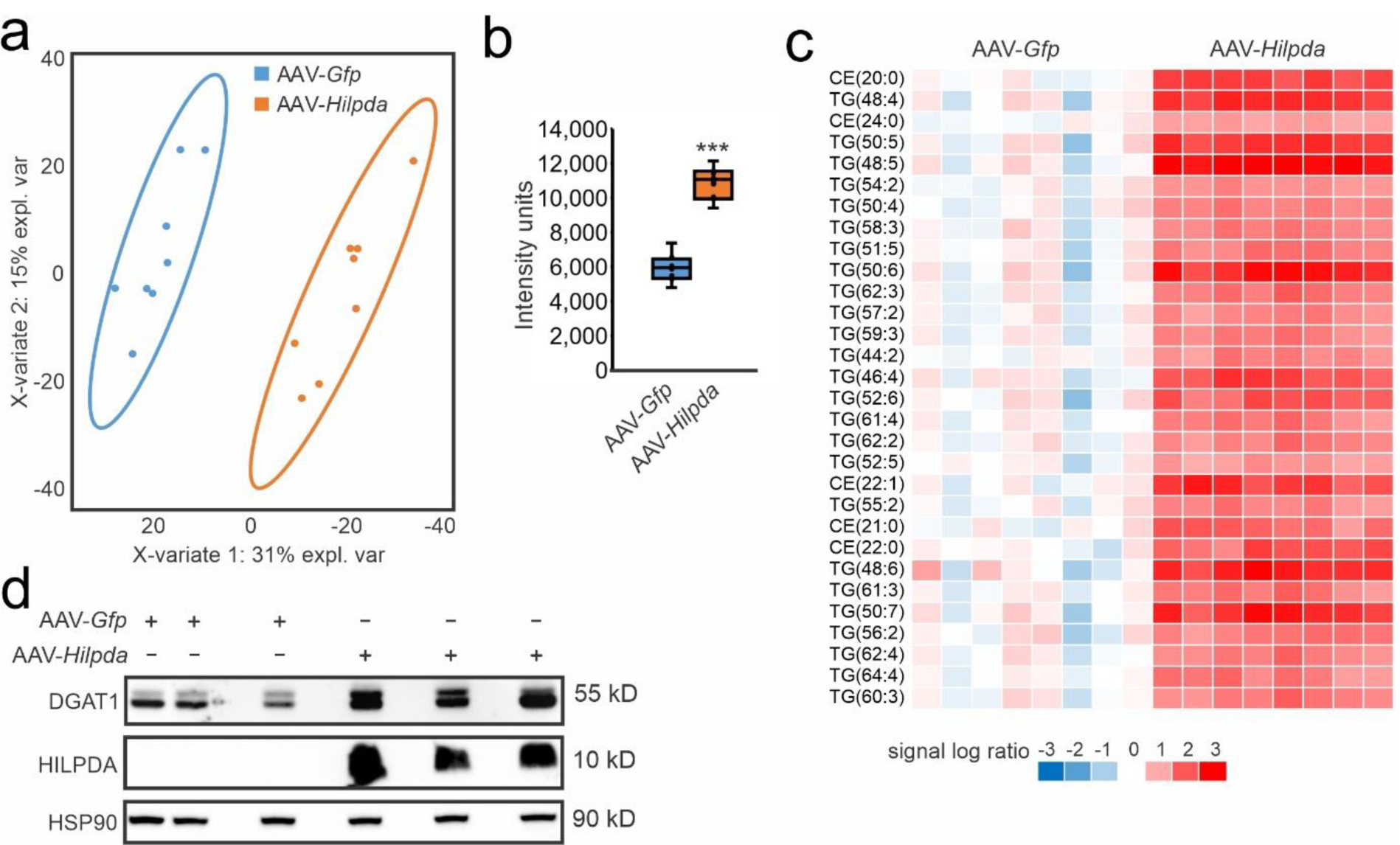
HILPDA overexpression promotes triglycerides storage and increases DGAT1 levels in mouse liver. Livers were collected from mice 4 weeks after injection with AAV-*Gfp* or AAV-*Hilpda* (24). a) PLS-DA analysis of the liver lipidomics profiles. b) Cumulative hepatic concentration of all triglyceride species. c) Heatmap of the 30 most significantly altered lipid species. d) DGAT1 and HILPDA protein levels in livers of mice infected with AAV-*Gfp* or AAV-*Hilpda*.

### HILPDA physically interacts with DGAT1

To investigate if HILPDA may physically interact with DGAT1 in cells, we performed FRET quantified by FLIM. In live HepG2 cells transfected with HILPDA-mEGFP and DGAT1-mCherry, HILPDA partially colocalized with DGAT1 (Figure 7a). Because confocal microscopy is diffraction limited to ∼250 nm, our colocalization results do not directly demonstrate that HILPDA and DGAT1 are physically interacting. To determine protein interactions, we performed FRET quantified by FLIM. The mean fluorescence lifetime of the donor fluorophore HILPDA-EGFP was significantly decreased by the presence of the acceptor fluorophore DGAT1-mCherry compared to the donor fluorescence lifetime in the absence of acceptor (Figure 7b-c). This result indicates that HILPDA and DGAT1 are in very close proximity, demonstrating a direct interaction between these two proteins. Transfection of HepG2 cells with HILPDA-mEGFP and DGAT2-mCherry showed that HILPDA also partially colocalizes with DGAT2 (Figure 7d). As for DGAT1, the mean fluorescence lifetime of the donor HILPDA-EGFP was significantly decreased upon co-transfection of the acceptor DGAT2-mCherry (Figure 7e-f) compared to the donor fluorescence lifetime in the absence of acceptor. These data demonstrate that HILPDA is able to physically interact with both DGAT1 and DGAT2. By contrast, although GPAT4-EFGP and HILPDA-mCherry showed substantial colocalization, FRET-FLIM analysis did not reveal a significant change in donor lifetime, indicating that these proteins do not interact (Supplemental figure 3a-c). Also, no significant change in donor lifetime was observed for HILPDA-mEGFP in combination with PLIN3-mCherry or GPAT1-EGFP in combination with HILPDA-mCherry (Supplemental figure 3d). Finally, HILPDA-mEGFP did not colocalize with PLIN2-mCherry (Figure 7g-h).

**Figure 7:**
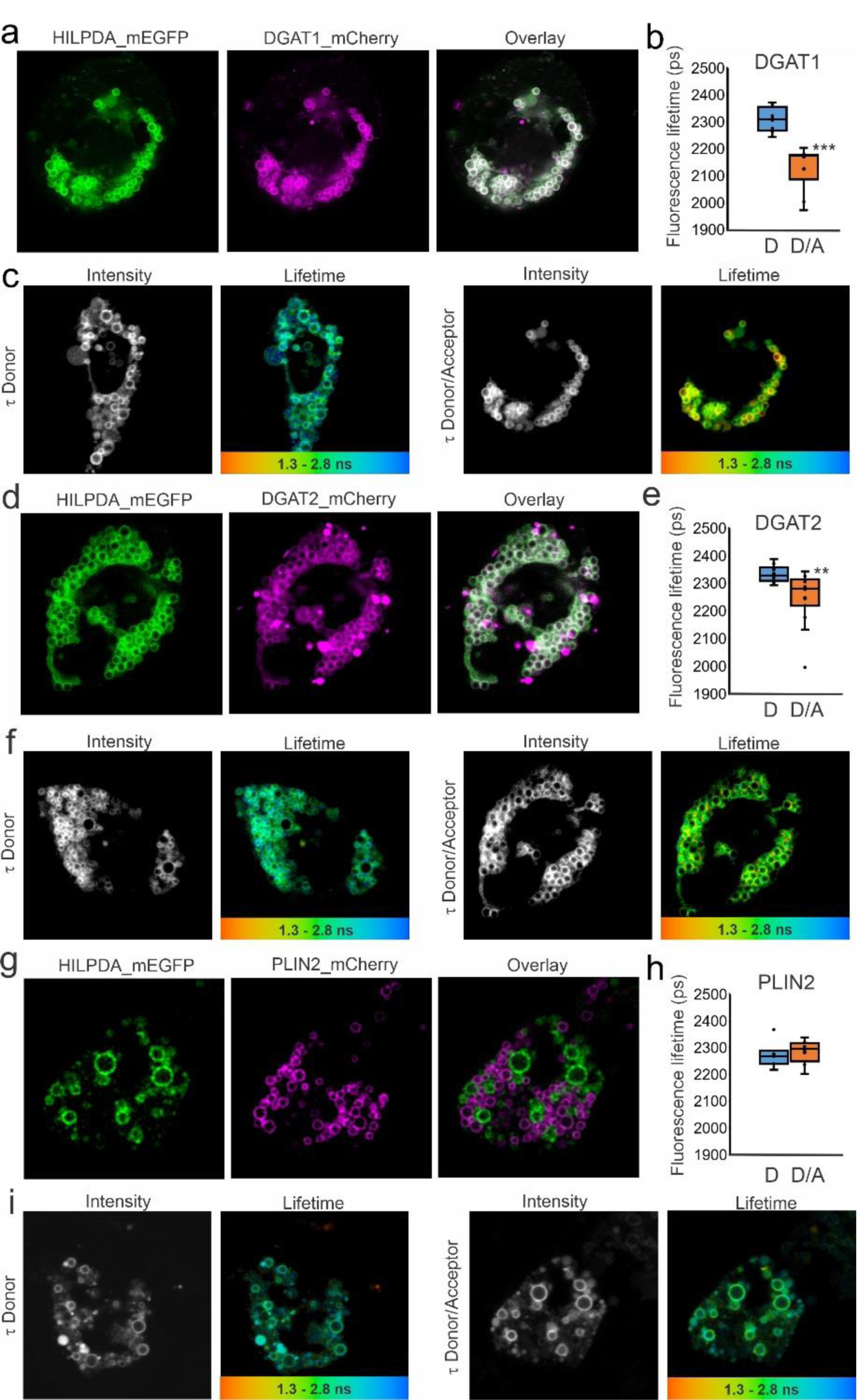
HILPDA and DGAT1/DGAT2 colocalize and physically interact intracellularly. HepG2 cells were transfected with HILPDA_mEGFP and DGAT1_mCherry or DGAT2_mCherry under lipid loaded conditions. Microscopy was carried out on live cells. a) HILPDA_EGFP and mDGAT1_mCherry partially colocalize in HepG2 cells. b) Fluorescence lifetime (τ) of HILPDA_EGFP in absence and presence of acceptor DGAT1_mCherry. c) Intensity image and LUT coloured lifetime image from red (1300 ps) to blue (2800 ps) from HILPDA_EGFP lifetime (τ) in the absence (left) or presence (right) of DGAT1_mCherry indicating where interaction occurs d) HILPDA_EGFP and DGAT2_mCherry partially colocalize in HepG2 cells. e) Fluorescence lifetime (τ) of HILPDA_EGFP in absence and presence of acceptor DGAT2_mCherry. f) Intensity image and LUT coloured lifetime image from red (1300 ps) to blue (2800 ps) from HILPDA_EGFP lifetime (τ) in the absence (left) or presence (right) of DGAT2_mCherry indicating where interaction occurs g) HILPDA_EGFP and PLIN2_mCherry do not colocalize in HepG2 cells. h) Fluorescence lifetime (τ) of HILPDA_EGFP in absence and presence of acceptor PLIN2_mCherry. Asterisk indicates significantly different from donor only according to Student’s t test; **P < 0.01; ***P < 0.001.

We repeated the FRET-FLIM experiments in fixed HepG2 and obtained similar outcomes. Specifically, co-expression of HILPDA-mEGFP with DGAT1-mCherry (Supplemental figure 4a-c) and DGAT2-mCherry (Supplemental figure 4d-f) led to a significant reduction in donor fluorescence lifetime. Collectively, these data indicate that HILPDA physically interacts with DGAT1 and DGAT2, but not with any of the other proteins studied.

## DISCUSSION

The purpose of this research was to better define the role and mechanism of action of HILPDA in liver cells. Consistent with previous studies (12, 24), we found that HILPDA stimulates lipid storage in hepatocytes. Interestingly, HILPDA mainly associated with active LD that are being remodelled. The stimulation of lipid storage by HILPDA appeared to be independent of ATGL and lipophagy. In HepG2 cells, HILPDA directly interacted with DGAT enzymes, stimulated DGAT activity, and increased DGAT1 protein levels. In vivo, the increase in hepatic triglyceride levels by HILPDA overexpression was accompanied by an increase in DGAT1 protein. Our data suggest that in addition to inhibiting ATGL-mediated lipolysis, HILPDA may increase intracellular lipid storage by stimulating triglyceride synthesis.

We and others previously showed that similar to the homologous G0S2 protein, HILPDA inhibits ATGL via a direct physical interaction, leading to suppression of triglyceride hydrolysis (28, 44). In macrophages and cancer cells, the stimulatory effect of HILPDA on lipid storage was almost entirely dependent on ATGL, suggesting that ATGL is the primary target of HILPDA in certain cell types (35, 36, 44). In adipocytes, physiological levels of HILPDA do not seem to have an impact on lipolysis, although at supra-physiological levels, HILPDA was able to reduce ATGL protein levels and inhibit lipolysis (11, 13). Interestingly, in hepatocytes, even though ATGL inhibition effectively increased lipid storage, the stimulatory effect of HILPDA on lipid storage was independent of ATGL. These data suggest that HILPDA not only inhibits ATGL but may also interact with other proteins involved in triglyceride turnover. These different interactions are likely cell type-specific. In fact, HILPDA may be part of a larger triglyceride turnover complex (“lipolysome”) that includes enzymes involved in triglycerides synthesis and triglyceride breakdown, including ATGL, as well as regulatory proteins such as ABHD5 and G0S2 (3, 9, 42).

As alluded to above, HILPDA and G0S2 share extensive homology, and both proteins are able to inhibit ATGL. Recently, evidence was provided that G0S2 not only suppresses lipolysis but also promotes triglyceride synthesis by carrying GPAT (LPAAT/AGPAT) enzymatic activity (45). Given the very small size of HILPDA (63 amino acids), it is unlikely that HILPDA can function as a fatty acid esterification enzyme. Rather, our data suggest that HILPDA serves as a small-protein activator of the DGAT1 enzyme. DGAT1 has been found to mediate esterification of exogenous fatty acids and fatty acids released from LD (8, 37). Interestingly, we observed that HILPDA localizes with active lipid droplets that are being lipolysed (disappear) and remodelled (form new LD). In addition, we found that HILPDA directly interacts with DGAT1, increases DGAT1 protein levels, and stimulates DGAT activity. Future studies will have to further clarify the precise molecular mechanism underlying the increase in DGAT1 protein by HILPDA overexpression.

In this paper, we show that deficiency of HILPDA in mouse liver led to a modest reduction in triglyceride storage after inducing NASH. Previous studies found that HILPDA deficiency did not significantly influence hepatic triglyceride levels in mice fed chow or a high fat diet (12). The reason for the divergent results is unclear but could be related to the different types of diets used. In any case, although statistically significant, the magnitude of the effect of HILPDA deficiency on hepatic triglyceride levels in mice was modest, which may be explained by the relatively low expression of *Hilpda* in mouse liver. By contrast, raising liver HILPDA levels by adeno-associated virus markedly elevates triglyceride storage (24).

Whereas HILPDA deficiency only had a modest effect on triglyceride storage in mouse liver, deficiency of HILPDA markedly reduced lipid storage in primary hepatocytes. This finding is consistent with the much higher *Hilpda* expression in primary hepatocytes compared to mouse liver. Accordingly, the effect of HILPDA deficiency and overexpression seem to depend on the baseline *Hilpda* expression. Indeed, specific physiological, pathological, and pharmacological stimuli may elevate HILPDA levels, thereby rendering HILPDA more important. A pathological condition associated with upregulation of *Hilpda* is infection with hepatitis C virus (19), which, interestingly, uses lipid droplets for replication (1, 27). Also, as HILPDA is highly induced by hypoxia, HILPDA is an excellent candidate to mediate the stimulatory effect of hypoxia/HIF1α on hepatic triglyceride levels, as seen in cancer cells (16, 44).

Our data showing reduced lipid droplet storage in HILPDA-deficient hepatocytes is consistent with the data by DiStefano and colleagues (12). According to their fatty acid flux data, the decrease in lipid storage is explained by a combination of decreased fatty acid uptake, increased fatty acid beta-oxidation, and increased triglyceride lipolysis. While triglyceride lipolysis is known to be directly targeted by HILPDA, it is unclear if fatty acid uptake and β-oxidation are as well, or if they are affected indirectly. Here we show that HILPDA also targets DGAT1-catalyzed triglyceride synthesis, supposedly via a direct interaction between HILPDA and DGAT1.

Currently, hardly anything is known about HILPDA in human liver. If the expression level of *HILPDA* in human liver is sufficiently high, inactivation of HILPDA could in theory be a promising strategy to treat non-alcoholic fatty liver disease. Whether NAFLD is associated with a change in the expression of *HILPDA* in human liver is unknown. Because loss-of-function variants in *HILPDA* would be expected to lead to reduced hepatic lipid storage, *HILPDA* is unlikely to emerge from any genome-wide association screens on NAFLD. Using multiple tools, we searched for SNP missense variants in the protein-coding region of the *HILPDA* gene. We identified several missense variants, a number of which was predicted to have a negative impact on protein structure. All of the identified missense variants are rare or very rare with MAF<0.1%. Accordingly, human genetic studies are unlikely to clarify the role of HILPDA in human liver.

As our gene targeting approach was directed towards HILPDA in hepatocytes, our conclusions are also limited to the role of HILPDA in these cells. The fact that albumin-Cre mediated HILPDA deletion only reduced hepatic *Hilpda* mRNA by about 50-60% suggests that *Hilpda* is expressed in other liver cell types as well, including possibly Kupffer cells, stellate cells, and endothelial cells. Given the important role of HILPDA in lipid storage in macrophages, it would be of interest to study the effect of LysM-Cre mediated HILPDA deletion on NASH and on lipid storage in Kupffer cells.

In our study, expression of HILPDA in liver cells was induced by fatty acids, which is consistent with the very sensitive upregulation of HILPDA by fatty acids in macrophages (23, 35). Besides activating *Hilpda* transcription via PPARs (23, 24), it is possible that fatty acids also specifically upregulate HILPDA at the protein level. The marked upregulation of *Hilpda* by fatty acids is likely part of a feed forward mechanism to properly dispose of the fatty acids by promoting their storage as triglycerides, either by activating the last step in triglyceride synthesis and/or inhibiting the first step in triglyceride breakdown.

Our study has several limitations. First, studies in cell culture were performed with overexpressed and tagged proteins, which may have influenced the results. It should be noted, though, that the colocalization of HILPDA to the ER and lipid droplets is fully in agreement with previous immunofluorescence studies on endogenous HILPDA(13, 23). Moreover, to study protein-proteins interactions in live cells via FRET-FLIM, it is necessary to overexpress and tag proteins. Second, the expression of *Hilpda* in mouse liver is low, certainly compared to macrophages, limiting the impact of HILPDA deficiency. Nevertheless, we could clearly detect HILPDA protein by Western blot in mouse liver, and observed a marked decrease in HILPDA abundance in hepatocyte-specific HILPDA-deficient mice. Third, direct evidence that physiological levels of HILPDA regulate triglyceride storage in mouse liver via DGAT1 is lacking. Addressing this question is technically extremely challenging. Instead, we showed that the stimulatory effect of HILPDA overexpression on triglyceride synthesis in liver cells is mediated by DGAT1.

In conclusion, our data suggest that HILPDA physically interacts with DGAT1 and increases DGAT activity in liver cells. Besides inhibiting ATGL-mediated lipolysis, HILPDA may increase lipid storage in cells by stimulating triglyceride synthesis.

## Acknowledgements

Funding from the Netherlands Organisation for Scientific Research (2014/12392/ALW), Consejo Nacional de Ciencia y Tecnologia de México (CONACYT) and from the Netherlands Cardiovascular Research Initiative (CVON2014-02 ENERGISE), an initiative with support of the Dutch Heart Foundation, is gratefully acknowledged. The authors would like to thank Matthijs Hesselink for valuable comments on the manuscript and Venetia Bazioti, Marialena Chrysanthou, Kaja Hribar, Anneke Hempel, Fabian Rood, and Shohreh Keshtkar for their help in carrying out the experiments,

## Notes

### Competing Interest Statement

The authors have declared no competing interest.

