## Supplemental figures for "Hypoxia-inducible lipid droplet-associated interacts with DGAT1 and promotes lipid storage in hepatocytes"

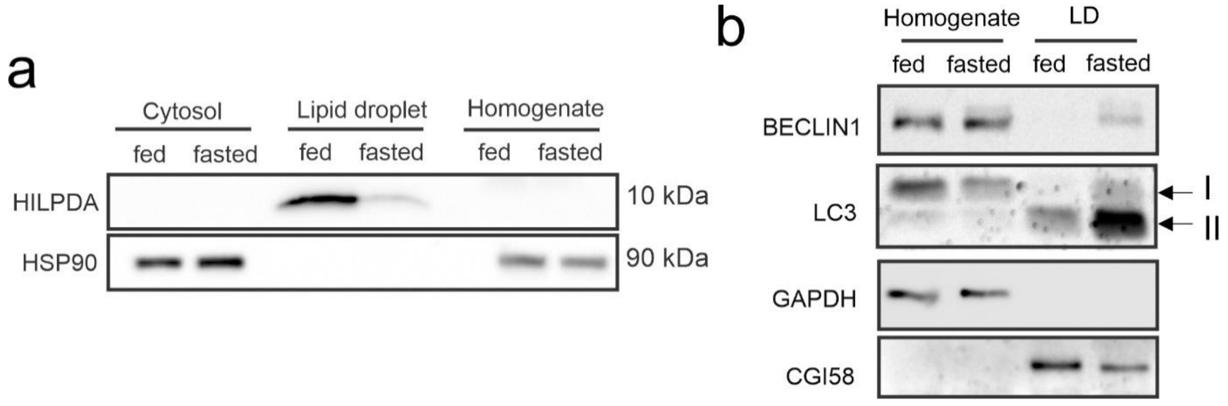


Supplemental figure 1. a) Enrichment of HILPDA in the lipid droplet fraction of fasted and fed wildtype livers as shown by immunblot. b) Immunoblots showing the presence of autophagy-related proteins Beclin 1 and LC in homogenate and lipid droplets fractions of livers of fed and 24h fasted mice. Five livers of wildtype C57Bl/6 mice were pooled into each sample. Absence of GAPDH validates absence of cytosolic contamination of lipid droplet fractions. Presence of CGI58 (ABHD5) demonstrates enrichment of the LD fraction.

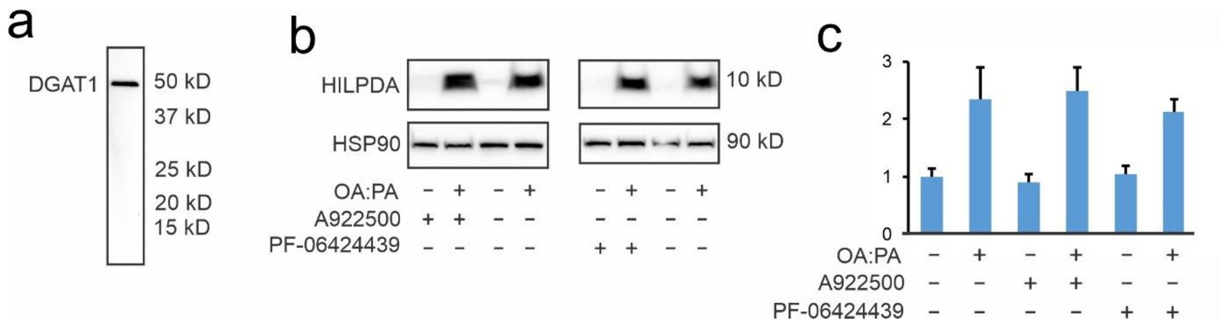


Supplemental figure 2. a) The DGAT1 antibody effectively detects DGAT1 in HepG2 cells transfected with DGAT1 expression vector. No effect of DGAT1 or DGAT2 inhibition on HILPDA protein levels (b) or mRNA levels (c) in Hepa1-6 cells. Hepa1-6 cells were pre-treated with DGAT1 inhibitor (A922500, 1 µM) or DGAT2 inhibitor (PF-06424439, 20 µM) for 30 minutes followed by co-treated for 24 hours with a 2:1 mixture of oleate and palmitate (total concentration (0.6 mM). Error bars represent SD.

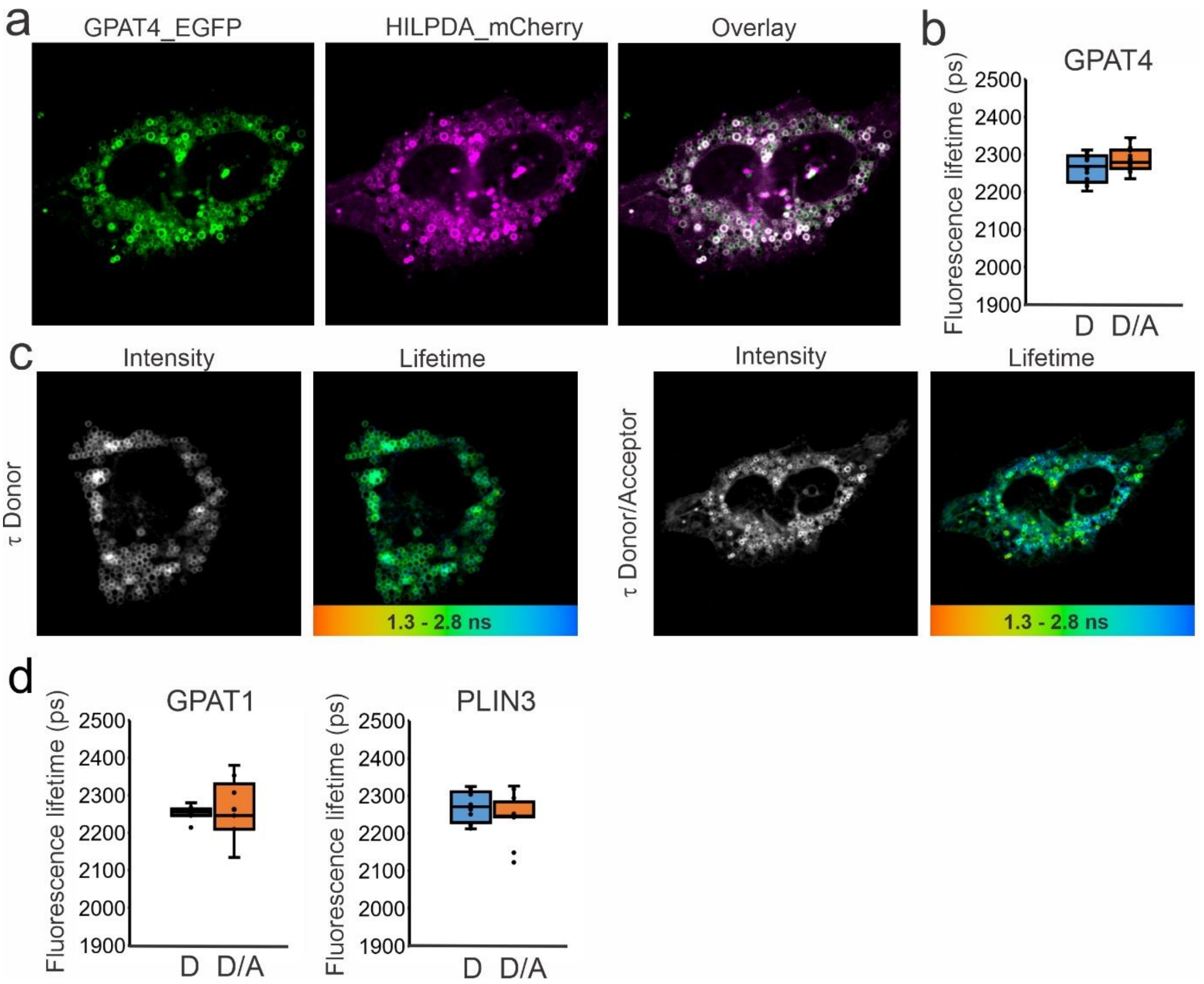


Supplemental figure 3. HepG2 cells were transfected with HILPDA_mCherry and GPAT4_EGFP, GPAT1_EGFP or HILPDA_EGFP and PLIN3_mCherry under lipid loaded conditions. Microscopy was carried out on live cells. a) HILPDA_mCheryy and GPAT4_EGFP colocalize in HepG2 cells. b) Fluorescence lifetime (τ) of GPAT4_EGFP in absence and presence of acceptor HILPDA_mCherry. c) Intensity image and LUT coloured lifetime image from GPAT4_EGFP lifetime (τ) in the absence (left) or presence (right) of HILPDA_mCherry. d) Fluorescence lifetime (τ) of GPAT1_EGFP in absence and presence of acceptor HILPDA_EGFP and fluorescence lifetime (τ) of donor HILPDA_EGFP in the absence and presence of acceptor PLIN3_mCherry.

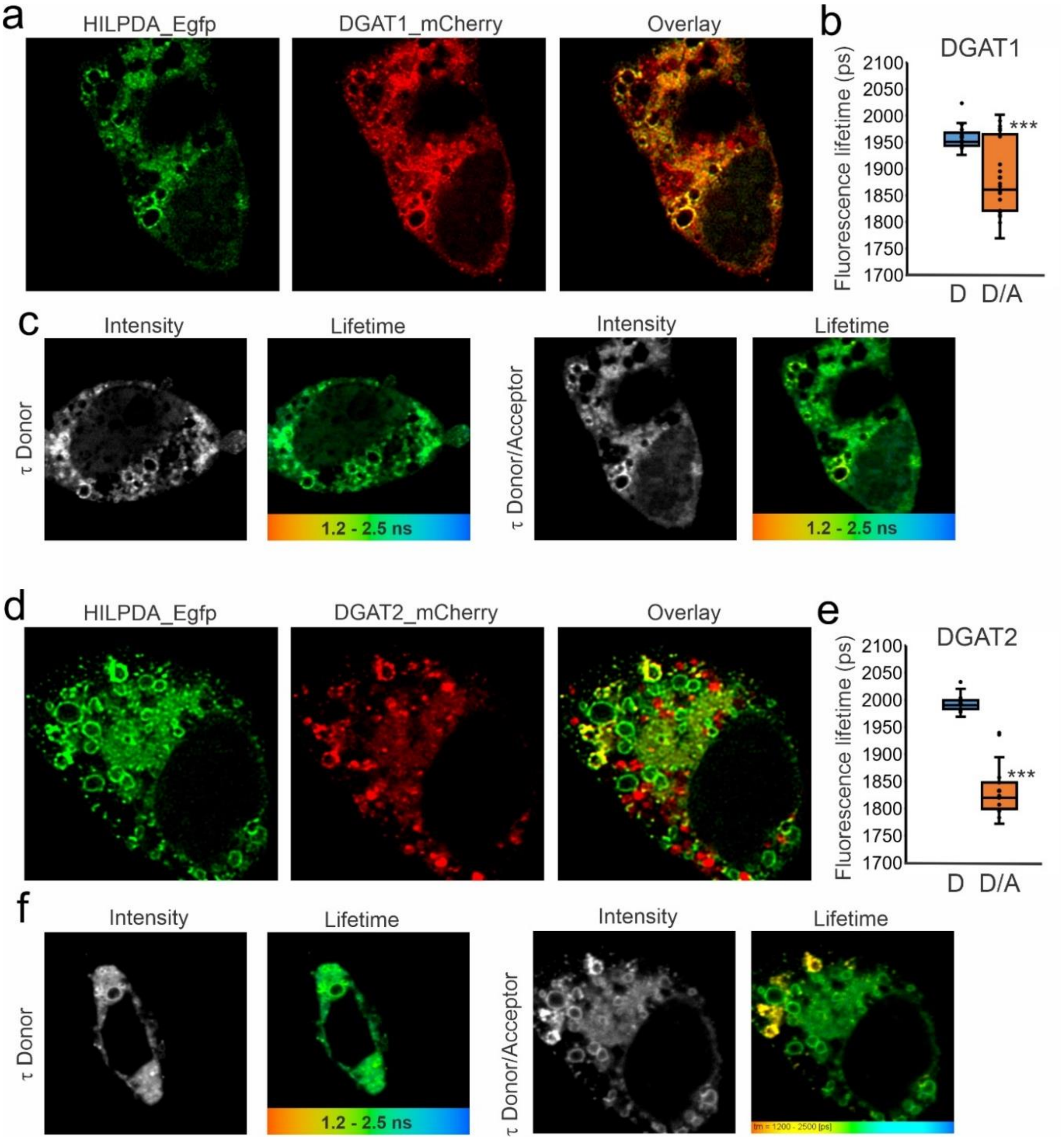


Supplemental figure 4: HILPDA and DGAT1/DGAT2 colocalize and physically interact intracellularly. HepG2 cells were transfected with HILPDA_EGFP and DGAT1_mCherry or DGAT2_mCherry under lipid loaded conditions. Microscopy was carried out on fixed cells. a) HILPDA_EGFP and mDGAT1_mCherry partially colocalize in HepG2 cells. b) Fluorescence lifetime (τ) of HILPDA_EGFP in absence and presence of acceptor DGAT1_mCherry. c) Intensity image and LUT coloured lifetime image from red (1200 ps) to blue (2500 ps) from HILPDA_EGFP lifetime (τ) in the absence (left) or presence (right) of DGAT1_mCherry indicating where interaction occurs) HILPDA_EGFP and DGAT2_mCherry partially colocalize in HepG2 cells. e) Fluorescence lifetime (τ) of HILPDA_EGFP in absence and presence of acceptor DGAT2_mCherry. f) Intensity image and LUT coloured lifetime image from red (1200 ps) to blue (2500 ps) from HILPDA_EGFP lifetime (τ) in the absence (left) or presence (right) of DGAT2_mCherry indicating where interaction occurs. Asterisk indicates significantly different from donor only according to Student’s t test; ***$P < 0.001$.
